# A novel microRNA-based strategy to expand the differentiation potency of stem cells

**DOI:** 10.1101/826446

**Authors:** María Salazar-Roa, Marianna Trakala, Mónica Álvarez-Fernández, Fátima Valdés-Mora, Cuiqing Zhong, Jaime Muñoz, Yang Yu, Timothy J. Peters, Osvaldo Graña, Rosa Serrano, Elisabet Zapatero-Solana, María Abad, María José Bueno, Marta Gómez de Cedrón, José Fernández-Piqueras, Manuel Serrano, María A. Blasco, Da-Zhi Wang, Susan J. Clark, Juan Carlos Izpisua-Belmonte, Sagrario Ortega, Marcos Malumbres

## Abstract

Full differentiation potential along with self-renewal capacity is a major property of pluripotent stem cells (PSCs). However, the differentiation capacity frequently decreases during expansion of PSCs in vitro. We show here that transient exposure to a single microRNA, expressed at early stages during normal development, improves the differentiation capacity of already-established murine and human PSCs. Short exposure to miR-203 in PSCs (*mi*PSCs) results in expanded differentiation potency as well as improved efficiency in stringent assays such as tetraploid complementation and human-mouse interspecies chimerism. Mechanistically, these effects are mediated by direct repression of *de novo* DNA methyltransferases Dnmt3a and Dnmt3b, leading to transient and reversible erasing of DNA methylation. As a proof of concept, miR-203 improves differentiation and maturation of PSCs into cardiomyocytes in vitro as well as cardiac regeneration in vivo, after cardiac injury. These data support the use of transient exposure to miR-203 as a general and single method to reset the epigenetic memory in PSCs, and improve their use in regenerative medicine.

## INTRODUCTION

Pluripotent stem cells (PSCs) provide an important promise for regenerative medicine due to their self-renewal potential and ability to differentiate into multiple cell lineages. During the last years, multiple efforts have been put into improving the potential of these cells either by expanding their differentiation capacity into a wider variety of cell lineages or by improving maturation properties into specific functional cell types^1, 2^. PSCs are commonly cultured in the presence of Mek1/2 and Gsk3 inhibitors together with the cytokine Lif (2i/L conditions; Ref. ^3^). Although these conditions may improve the maintenance of pluripotency in vitro, recent evidences suggest that prolonged inhibition of Mek1/2 may limit the developmental and differentiation capacity of PSCs in vivo, in part by inducing irreversible demethylation of imprinting control regions (ICRs)^4, 5^. Recent data suggest that genetic ablation of miR-34 in PSCs results in improved potential to form embryonic and extraembryonic tissues in part by promoting Gata2 expression^6^, although the conditions for applying this information to favor the differentiation potential of PSCs remain to be established. Finally, a recent alternative proposal suggests the use of a chemical cocktail of inhibitors that also enhances the developmental potential of PSCs^7^; however, the applicability of this method is still limited due to the lack of mechanistic details.

Here, we identified miR-203 as a microRNA preferentially expressed in the 2C-morula stages during preimplantation development in the mouse embryo. By using a variety of in vitro and in vivo approaches, we report that transient exposure of already-established induced pluripotent stem cells (iPSC) or ESCs to miR-203 enhances the ability of these PSCs to differentiate into multiple cell lineages and to reach further maturation properties. Transient expression of miR-203 in PSCs (*mi*PSCs) leads to greater developmental potential in tetraploid complementation assays, and significantly improves the efficiency of human iPSCs in contributing to interspecies human-mouse conceptuses. Mechanistically, these effects are at least partially mediated through the miR-203-dependent control of de novo DNA methyltransferases Dnmt3a and Dnmt3b, thereby regulating the DNA methylation landscape of these *mi*PSCs. These observations suggest that the developmental and differentiation potential of already established PSCs can be readily enhanced by transient exposure to a single microRNA.

## RESULTS

### miR-203 improves the differentiation potency of PSCs

miR-203 was originally proposed to limit the stemness potential of skin progenitors^8^ and to display tumor suppressive functions in multiple cancers^9, 10^, suggesting a role in the balance between stemness and differentiation. However, its expression during early development remained undefined. A first analysis of miR-203 levels during normal murine and bovine preimplantation development suggested a modest but specific wave of expression during the 2-cell stage and early blastocysts, whereas its expression was lost in cultured embryonic stem cells^11, 12^. Quantitative PCR analysis in mouse embryos isolated at different developmental stages showed that miR-203 expression was low in oocytes, slightly induced at the 2-cell stage and displayed a gradual reduction in morulas and blastocysts (Fig 1A).

**Figure 1.**
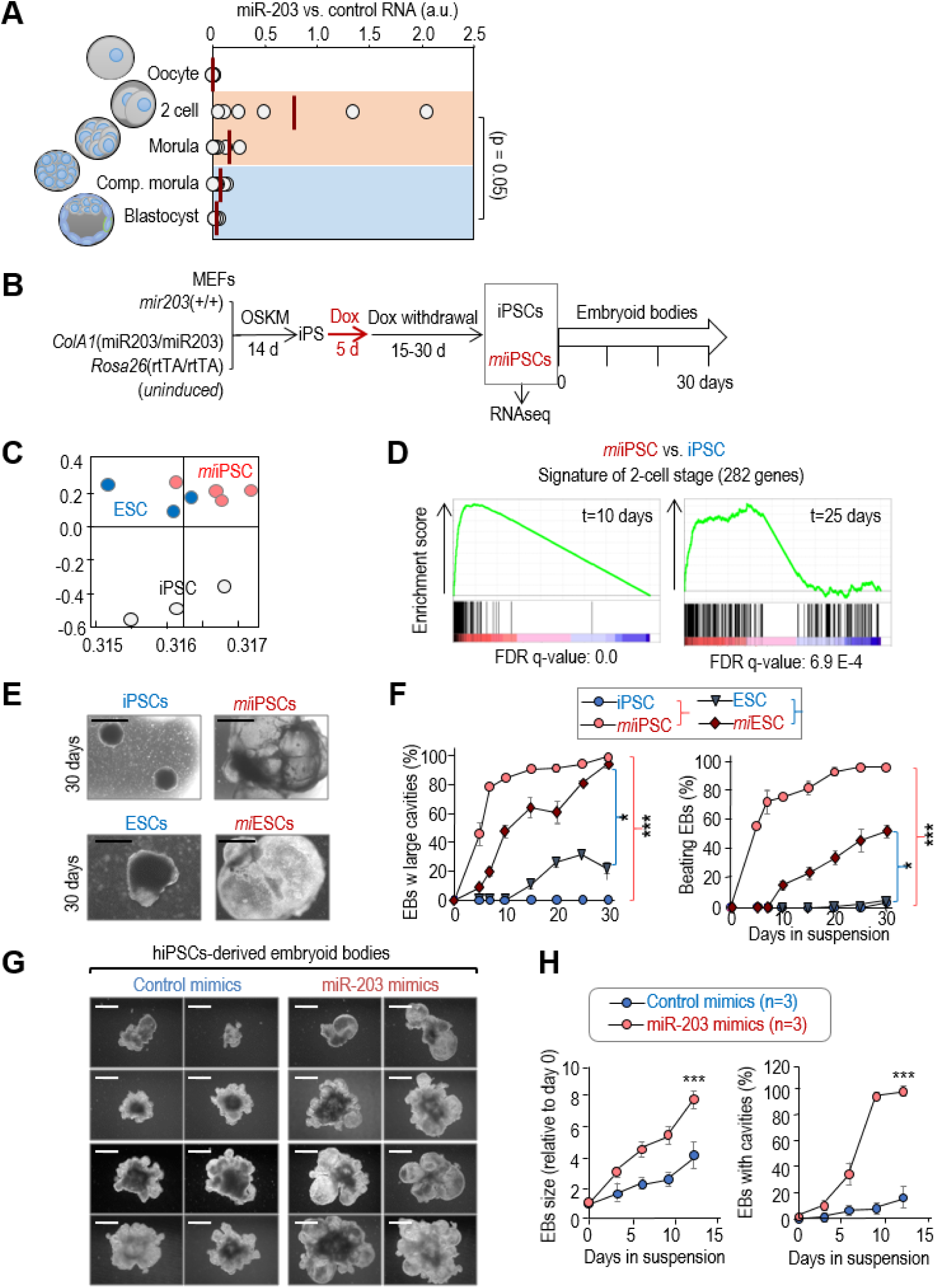
Effects of transient induction of miR-203 in iPSC and ESC pluripotency and differentiation potential. (**A**) miR-203 expression, as determined by qPCR, in five temporal different stages of normal early development: oocyte, 2-cell embryo, morula, compacted morula and blastocyst. RNA was extracted from 30 different embryos and pooled in two independent groups for analysis by qPCR. RNA expression is normalized by a housekeeping miRNA (miR-16) that maintained invariable during early embryogenesis. Data represent 6 different qPCR measures. *P*=0.05 (Student’s t-test) comparing 2C/morula versus compacted morula/blastocyst. (**B**) Protocol for reprogramming of miR-203 mutant MEFs into pluripotent iPSCs and subsequent differentiation into embryoid bodies. MEFs were transduced with lentiviruses expressing Oct4, Sox2, Klf4, and cMyc (OSKM) in a constitutive manner. The resulting iPSCs were then treated with doxycycline (Dox) 1 μg/ml during 5 days to induce miR-203 expression. “*mi*iPSCs” refers to iPSCs in which miR-203 was transiently expressed during the indicated 5 days. Dox was removed for 15-30 days before starting the embryoid body generation protocol. Samples for RNA sequencing were taken 30 days after Dox withdrawal. (**C**) Principal Component Analysis of RNAseq data from wild-type iPSCs (n=3 clones), *mi*iPSCs (n=4) and wild-type ESCs (n=3). (**D**) Enrichment plots of the 282-gene 2-cell signature^16^ in *mi*iPSCs 10 and 25 days after Dox withdrawal. (**E**) Representative images of embryoid bodies (EBs) derived from wild-type iPSCs or ESCs, or from *mi*iPSC and *mi*ESCs at day 30 of differentiation. Scale bars, 500 μm. (**F**) Quantification of the percentage of EBs from panel (**E)** presenting internal large cavities and EBs beating during the indicated time course. Data are represented as mean ± s.e.m. (n=3 independent experiments). \**P*<0.05; \*\*\**P*<0.001 (Student’s t-test). (**G**) Representative images of EBs derived from human iPSCs transiently transfected with either control (left) or miR-203 mimics (right), at different time points during the differentiation process. Scale bars, 500 μm. (**H**) Left panel shows the quantification of EBs size derived from human iPSCs transiently transfected with either control or miR-203 mimics as in panel (G) at different time points during the differentiation process. The percentage of EBs presenting internal large cavities during the indicated time course of differentiation is shown in the right panel. Data are mean ± s.e.m. (n=3 independent experiments). \*\*\**P*<0.001 (Student’s t-test).

To easily manipulating miR-203 levels in vitro and in vivo, we generated a tetracycline-inducible knock-in model in which the miR-203-encoding sequence was inserted downstream of the type I collagen gene and expressed under the control of a tetracycline-responsive element [*ColA1*(miR-203) allele] in the presence of tetracycline reverse transactivator expressed from the *Rosa26* locus [*Rosa26*(rtTA) allele] (Fig EV1A). Treatment of *ColA1*(miR-203/miR-203); *Rosa26*(rtTA/rtTA) mouse embryonic fibroblasts (MEFs) or ESCs with doxycycline (Dox) led to a significant induction of miR-203 levels (Fig EV1B). We also generated iPSCs from wild-type or un-induced *ColA1*(miR-203/miR-203); *Rosa26*(rtTA/rtTA) MEFs by using lentiviral vectors expressing Oct4, Sox2, Klf4 and cMyc (Ref. ^13^). Similar to MEFs or ESCs, these mutant iPSCs showed a significant induction of miR-203 after treatment with doxycycline (Fig EV1B). Although previous reports have suggested that this system suffers of relative leakiness^14^, miR-203 expression significantly reduced and was undetectable a few days after Dox withdrawal (Fig EV1C).

We next studied the long-term effects of miR-203 by treating *ColA1*(miR-203/miR-203); *Rosa26*(rtTA/rtTA) iPSCs (generated in the absence of Dox) transiently with Dox for 5 days (microRNA-transiently induced iPSCs or *mi*iPSCs; Fig 1B). RNA sequencing of these iPSC clones as well as wild-type ESCs was analyzed one month after miR-203 induction (Fig 1B) and surprisingly revealed that *mi*iPSCs cells were transcriptionally closer to ESCs than Dox-treated wild-type iPSCs both at the genome-wide level (Fig 1C) and when considering a pluripotency-associated signature defined previously^15^ (Fig EV1D). In addition, almost every transcript included in a 282-gene signature of 2C blastomeres^16^ was induced by miR-203 and sustained during ten days after Dox withdrawal (Fig 1D and EV1E), including genes such as Zscan4c previously related to the zygotic transcriptional program^17^. An additional comparison of the expression profiles in *mi*iPSCs versus un-induced iPSCs at day 30 after doxycycline withdrawal suggested higher expression of genes related to tissue morphogenesis and embryonic development in those clones in which miR-203 had been transiently induced (Fig 1B, EV1F and Supplementary Table 1).

The differentiation potential of *mi*iPSCs was directly tested in the embryoid body (EB) formation assay in vitro. Transient induction of miR-203 for 5 days followed by 2-4 weeks of Dox withdrawal in either iPSCs (*mi*iPSCs) or ESCs (*mi*ESCs) resulted in a significant increase in EB size accompanied by the formation of large internal cavities (Fig 1E,F). Furthermore, *mi*iPSC- and *mi*ESC-derived EBs showed beating with higher efficiency and at earlier time points than their untreated counterparts (Fig 1F). To further validate these observations beyond the genetic model, we tested the effect of transient expression of ectopic miR-203 using a CMV-driven retroviral vector or RNA mimics (Fig EV1G). These two strategies also resulted in improved EB formation (large cavities, beating) in *mi*iPSCs or *mi*ESCs (Fig EV1G), indicating that these effects were not unique to the inducible genetic model.

Finally, we decided to test the effects of miR-203 exposure on the differentiation potency of human PSCs. Thereby, we exposed hiPSCs to miR-203 mimics and analyzed their differentiation capacity in vitro, by embryoid bodies formation assays. Interestingly, human *mi*iPSCs were able to generate significantly bigger embryoid bodies with larger internal cavities than the control counterparts (Fig 1G,H), suggesting that the effect of miR-203 can be achieved using delivery systems with independence of the genetic knockin system and is also functional in human cells.

### miR-203 expands developmental potential and plasticity of PSCs

We then tested the potential of *mi*iPSCs (in which the expression of miR-203 had only been induced for 5 days in vitro) to form teratomas either after subcutaneous or intraperitoneal injection. *mi*iPSCs formed significantly bigger tumors in these assays (Fig 2A). Intriguingly, these teratomas were not only bigger but they contained tissues that are not typically found in control iPSC-induced teratomas, such as bone marrow, pancreas or cartilage, as well as trophoblast giant cells as confirmed by the expression of PL-1 (Fig 2B and Fig EV2). Transcriptomic studies in these complex structures suggested upregulation of genes involved in embryonic development and organ morphogenesis when derived from *mi*iPSCs (Supplementary Table 2).

**Figure 2.**
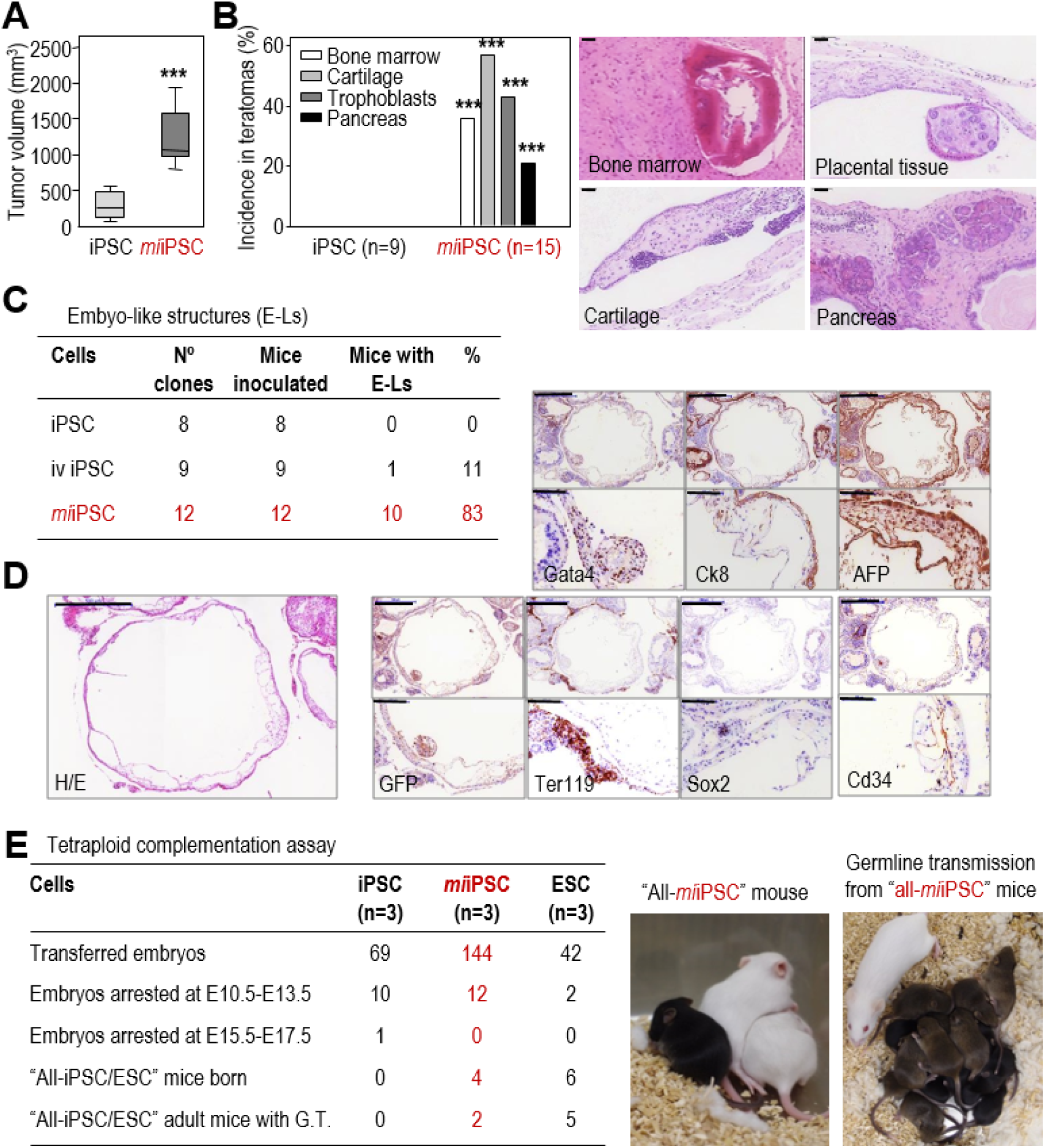
Transient exposure to miR-203 in vitro improves the in vivo developmental potential of iPSCs and ESCs. (**A**) Teratoma volume (mm^3^) 20-25 days after subcutaneous injection of wild-type iPSCs or *mi*iPSCs expressing GFP. Data are represented as mean ± s.e.m. (n=8 tumors per genotype); \*\*\**P*<0.001 (Student’s t-test). Representative images are shown in Fig EV2. (**B**) Incidence and representative images of specific highly differentiated tissues in teratomas. The number of tumors included in the analysis is indicated in the panel. \*\*\**P*<0.001 (Student’s t-test). Scale bars, 50 μm. (**C**) Table showing the frequency of nude mice with embryo-like structures (E-Ls) in their abdominal cavity 20-30 days after intraperitoneal (i.p.) injection of 400.000-500.000 in vivo (iv)-generated wild-type iPSCs, un-induced iPSCs, or *mi*iPSCs. The number of independent clones tested per condition is indicated in the panel (each animal was inoculated with a different clone). (**D**) Representative example of E-Ls generated after i.p. injection of GFP-expressing *mi*iPSCs. H&E, hematoxylin and eosin. The following antigens were detected by immunohistochemistry: GFP, Sox2 (ectoderm), Cd34 (mesoderm), Gata4 (endoderm), AFP and CK8 (visceral endoderm of the yolk sac) and Ter119 (nucleated erythroid cells). Scale bars, 500 μm and 100 μm for higher magnifications. (**E**) Embryo tetraploid complementation assays with un-induced iPSCs, *mi*iPSCs or wild-type ESCs (n=3 clones per condition). Pictures on the right show a representative example of a viable “all-*mi*iPSC” mouse (black) generated from *mi*iPSCs and litters obtained from “all-*mi*iPSC” adult mice, which efficiently contributed to germline transmission.

It has been reported that iPSCs generated in vivo are able to form small embryo-like structures, containing tissues derived from the three germinal layers in the presence of extraembryonic membranes, when inoculated intraperitoneally^18^. In our hands, wild-type iPSCs generated in vivo were able to form embryo-like structures in 11% of injected mice, with an efficiency similar to the one reported previously^18^. However, *mi*iPSCs generated in vitro were much more efficient, with 83% of injected mice showing embryo-like structures (Fig 2C). As in the case of in vivo-formed iPSCs, *mi*iPSC embryo-like structures were positive for specific markers of the three embryonic layers (Fig 2D).

The performance of *mi*iPSCs was also tested in the tetraploid complementation assay, one of the most stringent tests for developmental potency. Few iPSCs are efficient in this assay in which any born animal develops uniquely at the expense of exogenous diploid iPSCs (or ESCs) incorporated into a tetraploid embryo. Whereas we did not obtain all-iPSC mice in control iPSCs, the very same clones were able to form all-*mi*iPSC mice after transient in vitro exposure to miR-203 (Fig 2E). Importantly, half (2/4) of these “all-*mi*iPSC” mice reached adulthood healthy and were proficient for germline transmission (Fig 2E). Although specific clones of iPSCs have been previously found to contribute to all-iPSC mice^19^, it is important to consider that the effect of miR-203 is additive on already-established PSCs providing an easy protocol for enhancing available PSCs.

### miR-203 increases human-mouse interspecies chimera competency

The high efficiency of *mi*PSCs in contributing to mouse chimeras led us to test their effect in the formation of interspecies human-mouse conceptuses. Td-Tomato fluorescent-labeled human iPSCs were transiently transfected with pMCSV-miR-203-GFP (human *mi*iPSCs) and cells were maintained at least one week in culture to ensure that miR-203 expression was transient and absent before cell injection. To compare the effect of miR-203 in the context of the most efficient protocol reported so far, we cultured these human cells in the presence of human LIF, CHIR99021, (S)-(+)-Dimethindene maleate and Minocycline hydrochloride (LCDM), a cocktail recently proposed to enhance the potential of PSCs^7^. We injected one single human iPSCs into 8C-stage mouse embryos and examined its chimeric contribution after 48–60 h of culture in vitro. As shown in Fig 3A,B pre-exposure of human *mi*iPSCs to miR-203 resulted in 80-100% of the mouse blastocysts presenting Td-Tomato-positive human cells (also identified with anti-human OCT4 and HuNu antibodies), whereas this efficiency was only 50% when using control human iPSCs also cultured in the presence of LCDM but not miR-203. We next examined the contribution of human iPSCs to interspecies chimeras in similar assays in which injected 8C-stage embryos were allowed to implant. The presence of human cells in mouse E9.5 conceptuses was identified by immunostaining with anti-human nuclei (HuNu) antibody or by direct detection of fluorescent Td-Tomato, and their proper integration and differentiation into the mouse embryo was corroborated using embryo markers such as Gata4, Sox2 or hSOX17 (Fig EV3C). Both HuNu and Td-Tomato positive areas were significantly increased in embryos injected with *mi*iPSCs when compared to control iPSCs (Fig 3C-E and Fig EV3A,B). A similar effect of miR-203 was observed in chimera assays in which 15 Td-Tomato fluorescent-labeled human iPSCs, either control or transiently transfected with miR-203-expressing vectors (miiPSCs), were injected into 8C-stage mouse embryos and their contribution analyzed 48-60 h later (Fig EV3D), suggesting a more efficient contribution of human iPSCs to mouse embryos after brief exposure to miR-203.

**Figure 3.**
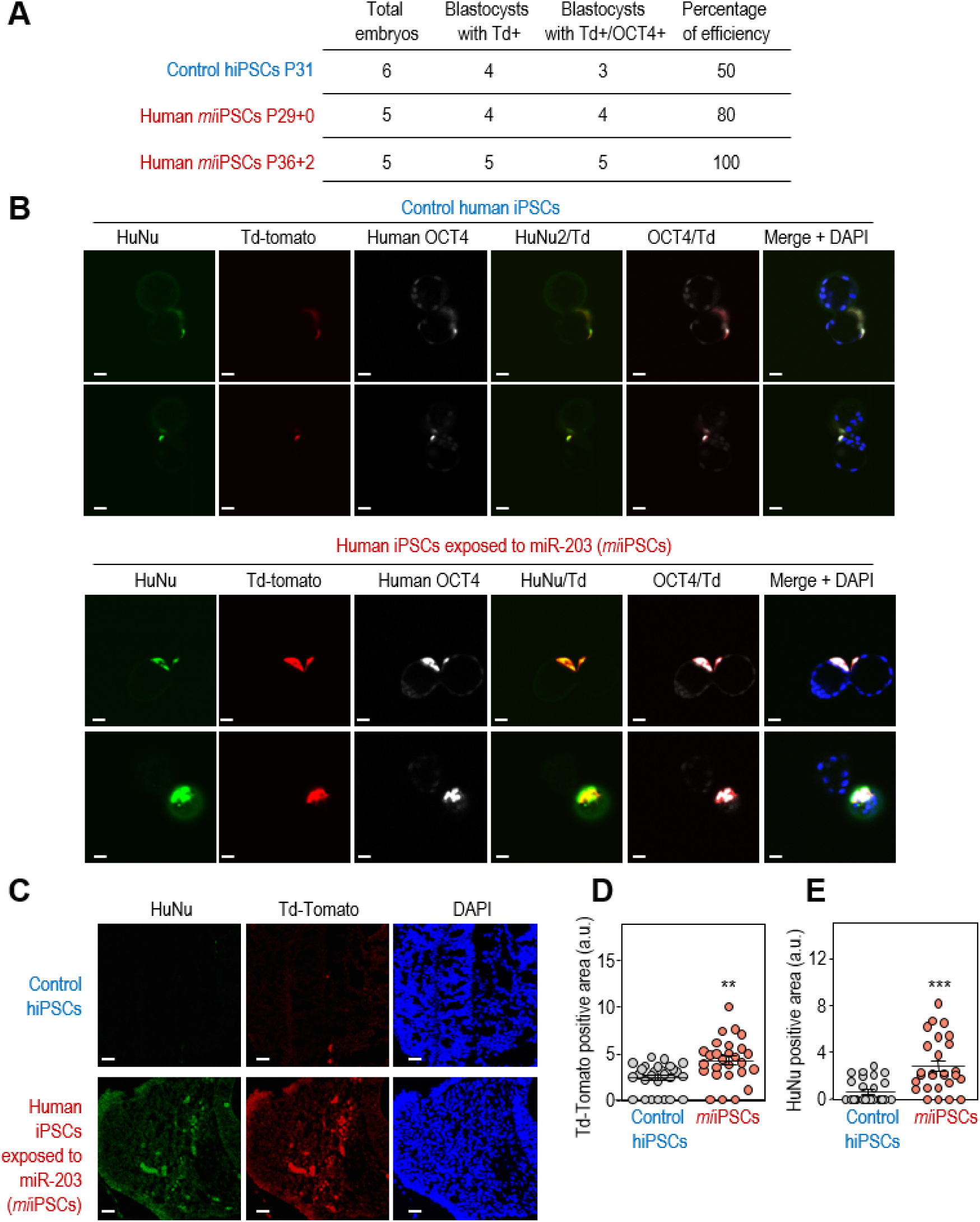
miR-203-exposed human iPSCs efficiently contribute to human-mouse interspecies chimeras. (**A)** Summary of chimera assays in which one single Td-Tomato fluorescent-labeled human iPSC, either control or transiently transfected with miR-203-expressing vectors (*mi*iPSCs) was injected into 8C-stage mouse embryos and their contribution analyzed 48-60 h later. (**B**) Representative images showing the contribution of human control iPSCs or *mi*iPSCs cells to mouse blastocysts. These structures were co-immunostained with (human) anti-OCT4 and anti-HuNu antibodies. Td-Tomato, red; Scale bars, 50 μm. (**C**) Interspecies chimera assays in which the 8C-stage mouse embryos indicated in (**A,B**), injected with one single human iPSCs and cultured in vitro for 48 h to reach the blastocyst stage, were then transfected to 2.5 days post coitum pseudo-pregnant females. The post-implantation mouse conceptuses were dissected at the E9.5 developmental stage and analyzed by immunofluorescence. Representative images showing the integration of control human iPSCs (top) or *mi*iPSCs (bottom) into mouse E9.5 embryos. Anti-human HuNu antibody was co-stained with Td-Tomato direct fluorescence to detect human cells in E9.5 mouse embryos. Scale bars, 20 μm. **(D,E**) Quantification of the Td-Tomato-(**D**) or HuNu-(**E**) positive area in 40-50 different cryosections of these embryos randomly and blindly selected for IF analysis. \*\**P*<0.01; \*\*\**P*< 0.001 (Student’s t-test).

### miR-203 effects are Dnmt3a/b-dependent

To analyze the mechanism by which miR-203 enhances the differentiation capacity of PSCs, we searched for predicted miR-203 targets among the transcripts downregulated in *mi*iPSCs (Fig 4A). As the long-term effects resulting from transient expression of miR-203 suggest an epigenetic mechanism, we selected from the previous list those genes involved in epigenetic regulation of gene expression (GO0040029). We found 18 GO0040029 transcripts downregulated in *mi*iPSCs cells and predicted as miR-203 targets according to different microRNA target prediction algorithms (Fig 4A and Supplementary Table 3). Among these transcripts, the top hits with the highest score as miR-203 targets corresponded to the *de novo* DNA methyltransferases *Dnmt3a* and *Dnmt3b.* These two transcripts were also significantly downregulated after expression of miR-203 mimics in wild-type iPSCs (log2 fold change= −0.24 for *Dnmt3a* and −0.22 for *Dnmt3b* transcripts). Human *miR*-203 and *DNMT3a/b* transcripts have been previously shown to display inverse expression profiles in cancer cells^20, 21^ and *DNMT3b* was recently shown to be a direct miR-203 target in human colon cancer cells^22^. Both *Dnmt3a* and *Dnmt3b* transcripts contain conserved miR-203 sites in their 3’-UTR (Fig EV4A) and exogenous expression of miR-203 led to decreased signal of a luciferase construct fused to these sequences, but not to 3’-UTR sequences from the related genes *Dnmt3l* or *Dnmt1* (Fig 4B,C). The downregulation of these genes was abrogated (Fig 4C) when the putative miR-203 binding sites were mutated (Fig EV4B) indicating a direct control of these transcripts by miR-203.

**Figure 4.**
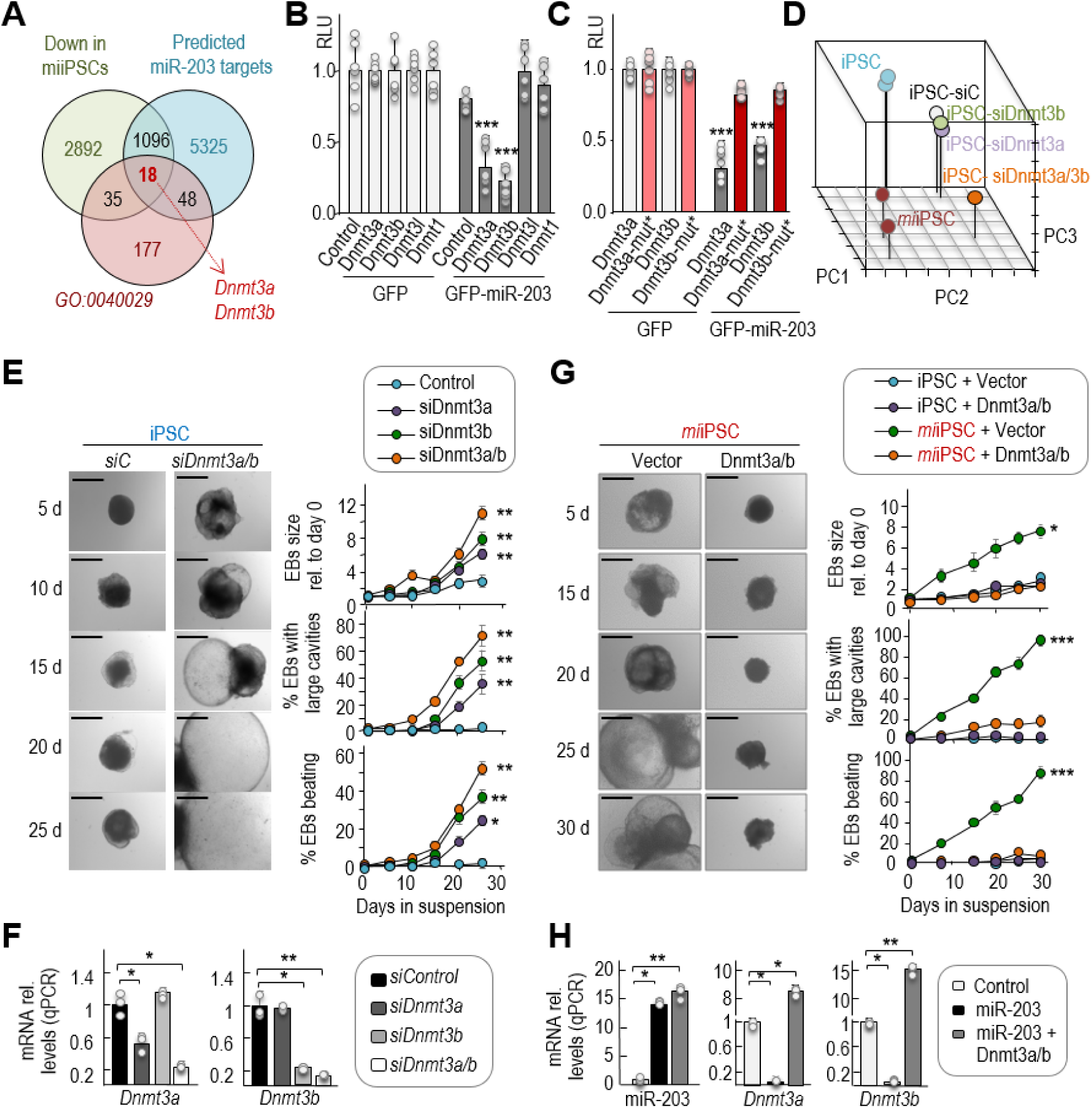
DNA methyltransferases 3a and 3b are miR-203 targets involved in the regulation of PSCs potential. (**A**) Venn Diagrams representing common genes down-regulated in *mi*iPSCs, predicted as miR-203 targets and also involved in the epigenetic regulation of gene transcription (GO:0040029). The list of the common 18 transcripts (including *Dnmt3a* and *Dnmt3b*) is presented as Supplementary Table 3. (**B,C**) Relative Luciferase Units (RLU; normalized to Renilla luciferase and relative to DNA amount) in 293T cells transfected with DNA constructs carrying the wild-type 3’UTRs from the indicated transcripts (**B**) or the mutated versions of *Dnmt3a* and *Dnmt3b* 3’UTRs, downstream of the luciferase reporter (**C**). Cells were co-transfected with *Renilla* luciferase as a control of transfection, and a plasmid expressing GFP or miR-203-GFP. Data are represented as mean ± s.d. (n=3 independent experiments). (**D**) Principal Component Analysis from RNAseq data including profiles from wild-type iPSCs, *mi*iPSCs, and wild-type iPSCs transfected with either control siRNAs (siC), or siRNAs specific against *Dnmt3a* (*siDnmt3a*), *Dnmt3b* (*siDnmt3b*) or both (*siDnmt3a*/b). (**E**) Representative images of embryoid bodies (EBs) derived from wild-type iPSCs in which the expression of *Dnmt3a* and *Dnmt3b* was transiently repressed by siRNAs. Scale bars, 500 μm. Plots show the quantification of the size of EBs and the percentage of EBs with large cavities or beating at different time points during the differentiation process. (**F**) Expression levels of *Dnmt3a* or *Dnmt3b* transcripts after transfection of wild-type iPSCs with specific siRNAs either against *Dnmt3a*, *Dnmt3b* or a combination of both (*Dnmt3a*/b). RNA expression was measured 24 hours after the transfection protocols and was normalized by GAPDH mRNA levels (n=3 independent experiments). (**G**) Representative images of EBs derived from *mi*iPSCs that were transiently and simultaneously transduced with Dnmt3a and Dnmt3b cDNAs or empty vectors, and simultaneously treated with Dox to induce miR-203 expression. Scale bars, 500 μm. Plots show the quantification of EB size, percentage of EBs with large cavities, and beating EBs at different time points during differentiation. (**H**) RNA expression was measured 24 hours after the transfection protocols and was normalized by a control miRNA (miR-142) or GAPDH mRNA, respectively. In (**B, C, E-H**) \**P*<0.05; \*\**P*<0.01; \*\*\**P*<0.001 (Student’s t-test).

To test to what extent downregulation of Dnmt3a/b could mimic the effect of miR-203 we knocked down these de novo DNA methyltransferases by RNA interference means (*siDnmt3a/b*). RNA sequencing of these samples (following the protocols explained above) revealed that *siDnmt3a*/b iPSCs exhibited a transcriptomic profile closer to *mi*iPSCs (Fig 4D and EV4C). In addition, downregulation of Dnmt3a/b induced the growth, formation of long cavities and increased beating of EBs to a similar extent to that observed in *mi*iPSC-derived EBs (Fig 4E,F), whereas individual knockdown of these transcripts displayed partial effects (Fig 4E,F and EV4D). Importantly, overexpression of miR-203-resistant Dnmt3a and Dnmt3b cDNAs, simultaneously to the transient doxycycline treatment, prevented the overgrowth of *mi*iPSC-derived EBs (Fig 4G,H and EV4D). Altogether, these observations suggest that Dnmt3a/b de novo methyltransferases are miR-203 targets whose downregulation facilitates the differentiation capacity of PSCs.

### miR-203 expression results in global and reversible DNA hypomethylation in PSCs

Given the critical role of Dnmt3a/b as *de novo* DNA methyltransferases we next analyzed the genome-wide methylation profile of control (un-induced) iPSCs and *mi*iPSCs (after transient induction with Dox) as well as embryoid bodies derived from them (Fig 5A). Wild-type and mutant iPSCs displayed similar levels of methylation before Dox and wild-type cells were not affected by this treatment. In contrast, transient induction of miR-203 for 5 days resulted in a genome-wide hypomethylation in *mi*iPSCs, which was strikingly more prominent 20 days after withdrawal of Dox (t=25; Fig 5B-E and Supplementary Tables 4-5), a time-point in which *Dnmt3a*/b transcript levels had already recovered after their repression in the presence of Dox (Fig EV5A). Notably, the number of DNA methylation valleys (DMVs; Ref. ^23^) and partially methylated domains (PMDs; Ref. ^24^) were increasingly higher in *mi*iPSCs with time (Fig 5B and Supplementary Table 5); for instance, 131 PMDs were found at t=0 while 548 and 6,555 PMD were found at t=10 and t=25, respectively. DNA methylation comparison between groups showed that 128 differentially methylated regions (DMRs) out of 131 total DMRs (97.7%; t=10 versus t=0) and 12,542 out of 12,549 (99.9%; t=25 versus t=0) DMRs were hypomethylated in *mi*iPSCs as a consequence of previous exposure to miR-203 (Fig 5E). These gradual changes in methylation spread over a significant proportion of Transcription Start Sites (TSSs) in a genome-wide manner (Fig 5F), and also affected genes of the 2C-signature (Fig 5G and Supplementary Table 6), in line with the observed upregulation in the expression levels of these 2C markers.

**Figure 5.**
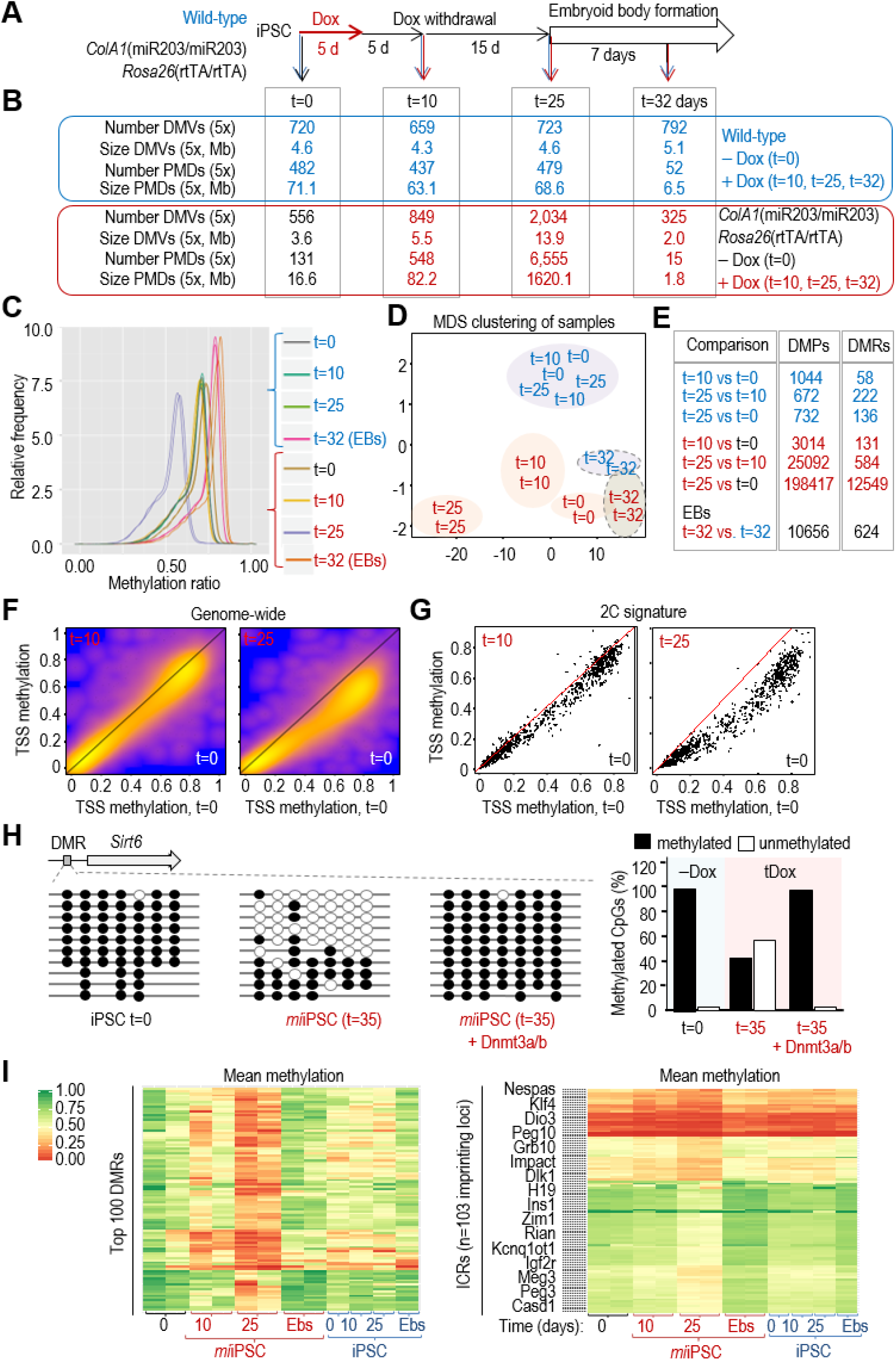
Transient expression of miR-203 induces genome-wide hypomethylation in iPSCs. (**A**) Experimental design for the genome-wide DNA methylation analysis of wild-type and *mi*iPSC as well as embryoid bodies (EBs) derived from them. Cells (two independent *mi*iPSC clones and two iPSC technical replicates) were transiently treated with Dox for 5 days and then subjected to Dox withdrawal for 20 additional days before starting the EB formation protocol. Samples for DNA and RNA analysis were collected at the indicated time points before Dox (t=0), 5 days after Dox withdrawal (t=10), 20 days after Dox withdrawal (t=25) or 7 days after starting the EB generation protocol (t=32 days). (**B**) Genome-wide DNA methylation data showing the number and total size of DNA methylation valleys (DMVs) and partially methylated domains (PMDs). (**C**) DNA methylation distribution of the indicated samples, smoothed over 100 kb-blocks. (**D**) Principal Component Analysis showing the distribution of methylation profiles in the indicated samples (color code as in panel **B**). (**E**) Number of differentially methylated single CpG sites (DMPs) and differentially methylated regions (DMRs) in the indicated comparisons (color code as in panel **B**). (**F**) Heat-scatterplots representing the mean methylation over all transcripts (transcription start sites, TSSs) in mm10, pairwise for “t=10 vs. t=0” (left panel) and “t=25 vs. t=0” (right panel). Note that especially for “t=25 vs. t=0” most of the points are below the diagonal, indicating broad hypomethylation across most of the genome. (**G**) Heat-scatterplots representing the mean methylation over the TSS of the 282 genes included in the 2-cell signature (as in Fig 1D). Analysis as in (**F**). (**H**) The specific differentially methylated regions (DMR) at the *Sirt6* locus was analyzed by PCR amplification and sequencing of bisulphite-modified DNA. The quantification of methylated vs. unmethylated CpGs is shown in the histogram in the indicated conditions. (**I**) Heatmaps representing the methylation levels at the top Differentially Methylated Regions (DMRs; n=100; left panel) or the Imprinting Control Regions (ICRs; n=103 different imprinting loci; right panel) in the indicated samples (referred to panels **A,B**). Color code and scale is applicable to both heatmaps.

As validation of the genome-wide methylation analysis, bisulphite modification followed by clonal sequencing confirmed miR-203-dependent hypomethylation of the promoter region of the gene encoding the E74 Like ETS Transcription Factor 5 gene (*Elf5*), a protein involved in the differentiation of epithelial cells and trophoblast stem-cell renewal and differentiation^25^ (Fig EV5B-E). Interestingly, the embryoid bodies derived from *mi*iPSCs displayed higher global levels of DNA methylation and lower number of PMDs (Fig 5B-E) in agreement with the upregulation of *Dnmt3a* and *Dnmt3b* transcripts observed after induction of differentiation (Fig EV5A). In line with the observations at a genome-wide scale, the *Elf5* promoter was hypermethylated in embryoid bodies generated from *mi*iPSCs during the differentiation process (Fig EV5B,C). Previous data suggest that interfering with DNA methyltransferase expression or activity results in global hypomethylation in the genome ^26–30^. Expression of exogenous, miR-203-resistant forms of Dnmt3a/b rescued the hypomethylation observed after miR-203 induction both in the *Elf5* DMRs (Fig EV5D,E) as well as in a DMR located at the histone deacetylase *Sirt6* gene (Fig 5H), suggesting that these de novo DNA methyltransferases are critical targets of miR-203 in inducing genome-wide hypomethylation.

Given the recent findings showing that a widespread loss of methylation in PSC cultures might be deleterious when accompanied by massive erasure of genomic imprints^4, 5^, we tested the methylation levels at 103 different genes controlled by Imprinting Control Regions (ICR) in *mi*iPSCs. Whereas *mi*iPSCs displayed a progressive demethylation of genes (red signal at t=10 and t=25 in Fig 5I; left panel), demethylation of ICRs was very limited at t=10 and moderate at t=25 in the same samples (right panel). Importantly, demethylation was in all cases fully recovered upon differentiation (*mi*iPSC-derived EBs; Fig 5I), suggesting that demethylation of *mi*iPSCs, both in DMRs and ICRs, is manageable and reversible, and does not compromise neither the quality of *mi*iPSCs nor their competence to differentiate.

### Transient exposure to miR-203 improves differentiation and maturation into cardiomyocytes as well as cardiac regeneration in vivo

Since transient expression of miR-203 improves the function of pluripotent cells in several assays, we next decided to directly testing the effect of expressing this miRNA during differentiation of PSCs into cardiomyocytes. The effect of miR-203 was first tested in primary cardiomyocytes isolated from neonatal (P1) rats that undergo further expansion and differentiation when cultured in vitro. miR-203 mimics triggered a transient burst of cell proliferation as measured by incorporation of the nucleotide analogue EdU (Fig EV6A) and mitotic markers such as cyclin B1 (Fig EV6B). Importantly, this increase in proliferation occurred in cardiac troponin T (cTnT)-positive cells (Fig EV6C) and resulted in cells with increased ratio of *Myh6* versus *Myh7* myosin heavy chain genes (Fig EV6B), a developmentally regulated switch that correlates with cardiomyocyte maturation and cardiac performance^31^.

We also tested a cardiomyocyte differentiation protocol from wild-type iPSCs^32^. Mouse iPSCs were transiently transfected with miR-203 or control mimics and 15 days later they were differentiated into cardiomyocytes using specific media and culture conditions (Fig 6A). Transient exposure to miR-203 was accompanied by increased expression of cardiomyocyte differentiation transcripts such as myosin heavy chain (*Myh*), atrial natriuretic peptide (*Nppa*), cardiac troponin T (encoded by the *Tnnt2* gene), markers for cardiac progenitors such as insulin gene enhancer transcription factor *Isl1* and *Tbx5* in iPSCs previously treated with miR-203 mimics (Fig 6B and EV6D). Importantly, transient exposure to miR-203 resulted in higher expression of not only differentiation but also maturation markers such as the potassium channel components encoded by the *Kcnh2*, *Hcn1* and *Kcna4* genes^33^, that were only minimally induced in cells treated with control mimic RNAs (Fig EV6E). In line with these data, the beating frequency in cardiomyocytes derived from miR-203-treated iPSCs was significantly higher suggesting enhanced functionality (Fig 6C). Expression of miR-203-resistant forms of Dnmt3a and Dnmt3b in parallel to miR-203 (Fig 6A) prevented the upregulation of these differentiation and maturation markers (Fig 6B,D) as well the increase in beating frequency (Fig 6C), suggesting the relevance of the miR-203-Dnmt3a/b axis in functional differentiation and maturation of cardiomyocytes from pluripotent cells.

**Figure 6.**
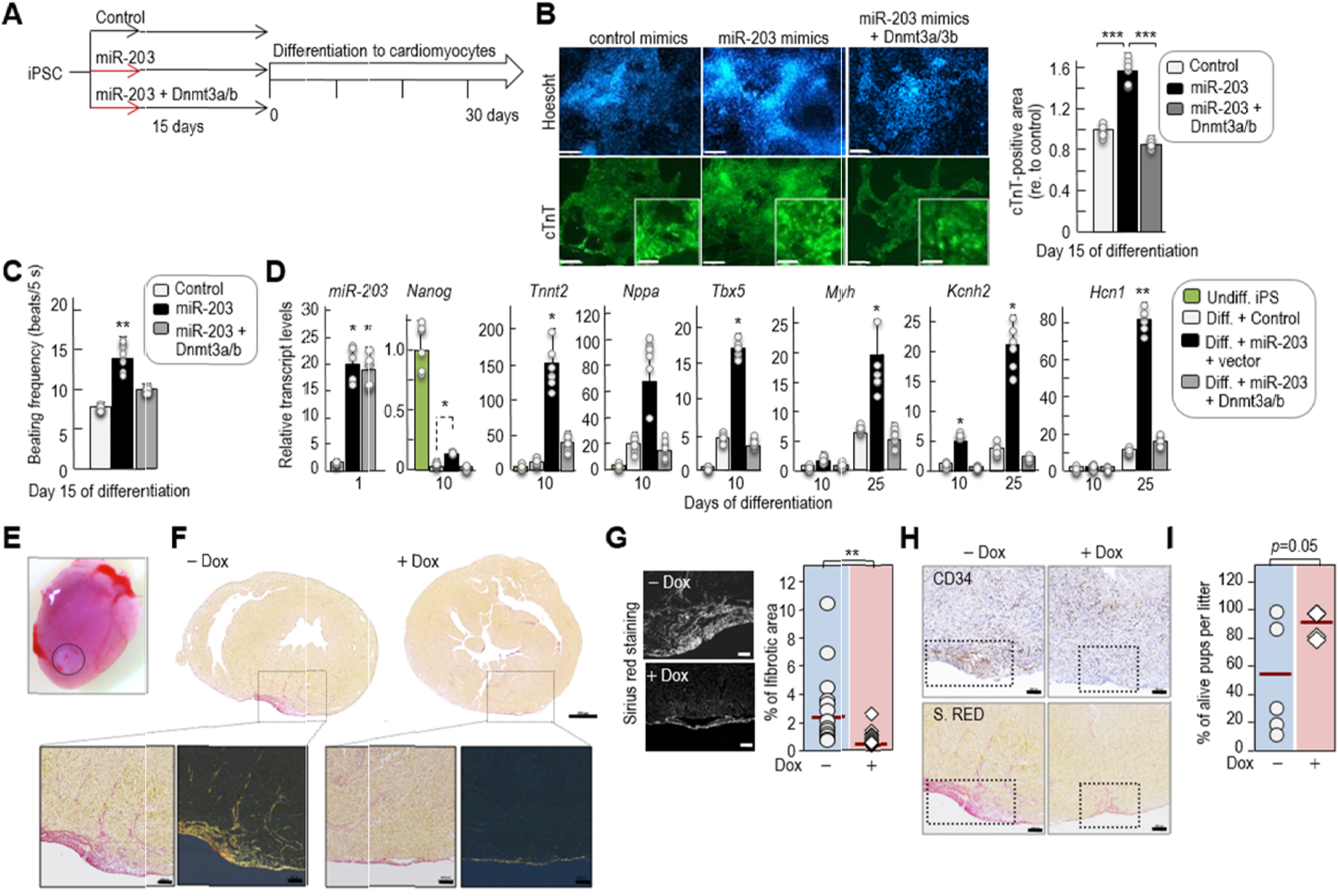
Transient exposure to miR-203 enhances differentiation into mature cardiomyocytes and improves cardiac regeneration. (**A**) Experimental protocol followed for the differentiation of cardiomyocytes from iPSCs in the absence or presence of miR-203 mimics and Dnmt3a/b cDNAs. (**B**) Representative immunofluorescences showing cardiac Troponin T (cTnT, green) and nuclei (Hoescht, blue) staining of in vitro-generated cardiomyocytes derived from wild-type iPSCs transiently transfected with either control mimics, miR-203 mimics or miR-203 mimics + Dnmt3a/b cDNAs. Pictures were taken at day 15 of differentiation. Lower panels show a magnification detail of cTnT staining in each condition. Scale bars, 68 μm (inset, 25 μm). The cTnT-positive area in these cardiomyocytes is shown in the right histogram. Data are represented as mean ± s.d. (n=2 independent experiments with 6 replicates each). (**C**) Beating frequency (measured as number of beats per 5 seconds) of these cardiomyocytes at day 15 of differentiation. (**D**) miRNA or mRNA levels as determined by quantitative PCR of the indicated transcripts at different time points during cardiomyocyte differentiation in the indicated samples. Data are represented as mean ± s.d. (n=2 independent experiments with 6 replicates each). (**E**) Representative image of a postnatal-day-8 heart, 7 days after the cryoinjury. The area of cryolesion is highlighted. (**F**) Representative images of heart sections from vehicle and Dox-treated mice stained with Sirius Red (scale bars, 500 μm). The magnification details show the fibrotic area in white (left) and polarized (right) light. Scale bars of the insets, 100 μm. (**G**) Representative “black and white” images of the heart sections stained with Sirius Red from representative vehicle- or Dox-treated mice, showing in white the fibrotic area 7 days after the cryoinjury. Scale bar, 250 μm. The histogram shows the quantification of the percentage of fibrotic area relative to the total heart area in n=15 mice per condition, 7 days after the cryoinjury. \*\**P*<0.01 (Student’s *t*-test). (**H**) Immunohistochemical detection of Cd34 (mesodermal progenitors) and Sirius Red staining of heart sections from vehicle and Dox-treated mice. Representative images from 3 different mice are shown. Scale bars, 100 μm. The fibrotic area is highlighted. (**I**) Quantification of the percentage of living pups per litter one day after the cryolesion. 5 litters were tested per condition. P= 0.053 (Student’s *t*-test). In (**B, C, D, G**), *\*P*<0.05; *\*\*P*<0.01; *\*\*\*P*<0.001 (Student’s *t*-test).

The fact that miR-203 increases cell plasticity to differentiate, inspired us to test the effect of miR-203 in cardiac regeneration after injury. The neonatal mouse cardiac cryoinjury enables testing of molecular interventions to stimulate regeneration, in a model with characteristics similar to those observed in pediatric heart disease^34^. Heart muscle cell death was induced by cryoinjury in *ColA1*(miR-203/miR-203); *Rosa26*(rtTA/rtTA) mice at postnatal day 1 (Fig 6E), and neonates were treated with vehicle (control) or Dox (to induce miR-203 expression) for seven days. After 1 week of recovery, control pups displayed significantly larger fibrotic areas in the heart, whereas wound healing was significantly improved in Dox-treated neonates (Fig 6F,G). Identification of CD34-positive cardiac progenitors in parallel to Sirius Red staining revealed an accumulation of undifferentiated progenitors in the scar of control hearts. Treatment with doxycycline, however, led to a significant reduction of fibrosis accompanied by reduced presence of these progenitors in the injured area and a better recovery of normal tissue (Fig 6H). Of interest, the percentage of living pups in the operated litters was higher in the Dox-treated group compared to controls (Fig 6I), suggesting the therapeutic effect of inducing miR-203 during cardiac regeneration.

## DISCUSSION

Multiple efforts in the last decade have focused on improving the developmental and differentiation potential of PSCs with the ultimate goal of using these cells in regenerative medicine (Supplementary Table 7). PSCs, and in particular iPSCs, are inefficient in the most stringent assays such as tetraploid complementation^35, 36^ and human-mouse interspecies assays^37, 38^. Improving culture conditions has been a major subject of research offering multiple options^4, 5, 7^. The use of 2i/L clearly improves some properties of PSCs in vitro and has been widely used in the last years, although recent data indicates that prolonged use of these conditions results in reduced developmental potential in vivo^4, 5^. Recent alternatives include the use of Src inhibitors or the LCDM cocktail whose targets require further validation^4, 5, 7^. Despite these advances, the long-term effects of chemicals that perturb major signaling pathways in the cell remain uncertain. In particular, it will be critical to study how these treatments affect the differentiation potential of PSCs towards specific cell lineages and the functional properties of the resulting cells.

In this work, we report that transient expression of a single microRNA can expand functionality and differentiation plasticity of either mouse or human PSCs both in vitro and in vivo. Already-established PSCs display enhanced expression of a signature of the top ∼280 transcripts characteristic of the 2-cell stage during embryonic development^16^ after brief exposure to miR-203. Importantly, transient exposure to this microRNA results in *i*) enhanced differentiation into multiple tissues (e.g. including pancreas, bone marrow or trophoblast) in teratomas generated in vivo; *ii*) complex embryo-like structures observed after the injection of these cells into mice; *iii*) augmented efficiency in stringent tests for assessing developmental potency, such as tetraploid complementation assays and human-mouse interspecies chimeras. Remarkably, half of those all-iPSC mice produced with *mi*iPSCs by tetraploid complementation reached the adulthood showing efficient germline transmission, which has been rarely found in previous reports^4, 5, 7, 35, 36, 39–41^. Similarly, human iPSCs cells transiently exposed to miR-203 contributed to interspecies human-mouse conceptuses with high efficiency (up to 100% contribution in vitro) compared to previous reports^37, 38, 42^. Importantly, miR-203 was able to enhance the potential of PSCs even when cultured in optimal culture conditions leading to the best in vitro-pluripotent cells (EPS) reported so far^7^, suggesting that exposure to this microRNA may be used in combination with other strategies aimed to optimize PSC culture conditions.

Endogenous miR-203 levels are transiently induced during early preimplantation development, suggesting that this microRNA may contribute to the specific expression profile of early blastomeres. The fact that brief exposure to miR-203 sequences improves the long-term developmental potential of *mi*PSCs even many weeks and passages after its expression suggests an epigenetic mechanism. In fact, *mi*PSCs display a hypomethylated state in chromatin that is self-sustained and accentuated several weeks after exposure to this microRNA implying a replication memory. Among the multiple miR-203 targets, de novo DNA methyltransferases Dnmt3a and Dnmt3b likely play an important role in this effect. In cancer cells, miR-203 expression has been previously shown to decrease DNMT3B protein levels leading to de novo methylation of cancer-related genes and decrease expression of their products including the xenobiotic transporter ABCG2^21, 22^. In ESCs, lack of these DNA methyltransferases is known to induce progressive hypomethylation of chromatin with passages^26, 28^ and knockdown of both *Dnmt3a and Dnmt3b* transcripts mimicked the effect of miR-203 in several assays. In addition, expression of miR-203-resistant-Dnmt3a and -Dnmt3b cDNAs rescue the phenotypes induced by miR-203.

Of note, whereas the severe hypomethylation observed in *Dnmt3a*/b knockout cells blocks differentiation^30, 43^, the hypomethylated state induced by miR-203 is reversible and differentiation is highly improved, when compared to untreated cells, and accompanied of potent de novo DNA methylation. Whereas the basis for these differences is unclear, given the promiscuity of microRNAs, miR-203 may have additional targets that could also cooperate with *Dnmt3a*/b repression in the effects of this microRNA in pluripotent cells.

Maintaining methylation of imprinted loci during preimplantation development is a process mediated by Dnmt1, rather than Dnmt3a or Dnmt3b^44, 45^, in agreement with our data showing a limited effect in the demethylation of ICRs in *mi*PSCs, in which Dnmt1 expression is not modified. Conditions in which ICRs are also demethylated are not appropriate for PSC function as ICR hypomethylation is frequently irreversible^42^. In fact, the use of 2i/L conditions result in a widespread loss of DNA methylation including massive erasure of genomic imprints that do not recover methylation after differentiation protocols^5777^, leading to defective developmental potential in chimerism studies or tetraploid complementation assays^4, 5^. On the contrary, transient exposure to miR-203 induces moderate and reversible ICR hypomethylation, that likely contributes to the differentiation and development potential of these PSCs when submitted to differentiation stimuli.

The developmental potential of *mi*PSCs can be also exploited in differentiation protocols towards specific cell types. As an example, here we have shown how exposure to miR-203 improves not only differentiation of PSCs into cardiomyocytes but also increases the expression of maturation markers such as components of the potassium channels involved in heart rate, and enhances contractibility of the resulting cells. Importantly, our genetic model allowed us to demonstrate the positive effect of miR-203 induction during tissue regeneration in vivo, using a cryoinjury model of relevance not only in translational heart failure research but also in pediatric-specific disease mechanisms and therapies^46^.

Altogether, our data present miR-203 sequences as an easy-to-use and suitable tool to improve PSCs in vitro, by expanding their differentiation and developmental potential in vivo. In contrast to other protocols aimed to improve reprogramming into pluripotent cells^1, 2, 29, 47, 48^, the effects induced by miR-203 can be easily achieved through brief exposure to specific mimics in already-established pluripotent cultures. Together, we conclude that transiently modulating the DNA methylation landscape of *mi*PSCs by short exposure to miR-203 may enhance the performance of these cells in multiple functional assays, including the efficient differentiation and maturation to specific cellular lineages of interest in regenerative medicine.

## ON-LINE METHODS

### Animal models and procedures

Animal experimentation was performed according to protocols approved by the CNIO-ISCIII Ethics Committee for Research and Animal Welfare (CEIyBA). The miR-203 inducible model was generated by cloning a 482-bp genomic *mmu-mir203* sequence into the pBS31 vector for recombination into the *ColA1* locus in ES cells following the strategy reported previously^49^. The resulting knockin allele [*ColA1*(miR-203)] was combined with a Rosa26-M2rtTA allele [*Rosa26*(rtTA)] for doxycycline-dependent induction as described previously^49^ (Fig EV1A). These animals were maintained in a mixed C57BL6/J x 129 x CD1 genetic background. For subcutaneous teratomas, iPSCs were trypsinized and 2-3 million cells were suspended in 100 μl PBS supplemented with 0.1% glucose and were subcutaneously injected into both flanks of athymic nude mice (provided by Charles River). Teratomas were daily controlled and measured by a caliper and finally isolated when their diameters reached 1.5 cm. Animals were euthanized at that point and teratomas were weighted and processed for RNA extraction or histopathological analysis. For intraperitoneal injections, wild-type athymic mice were injected with 4-5 x 10^5^ iPSCs resuspended in 100 μl PBS supplemented with 0.1% glucose. Mice were supervised daily from the day of the injection. Usually, 30 or 40 days after injection mice were euthanized and the visceral teratomas and embryo-like structures were isolated and processed, either for RNA extraction or histopathological analysis.

For tetraploid complementation studies, 2-cell stage Hsd:ICR(CD-1) embryos were harvested from pregnant females at E1.5 and electrofused in 0.3 M mannitol using a BLS CF-150/B cell fusion instrument with a 250 μm electrode chamber. The electric pulse conditions were 30 V amplitude, 31 μs width and 1.5 AC voltage. One to two hours later, 1-cell (tetraploid) embryos were selectively picked and cultured in KSOM overnight. Next day, 4-cell stage embryos were selected and aggregated with ESCs or iPSCs. Aggregated embryos were transferred to pseudo-pregnant females 24 hours later. To study germline contribution, black ES-iPS-mice (from tetraploid complementation assays) or chimeras (from diploid embryo ES-iPS microinjection) were crossed with Albino C57BL/6J-Tyrc-2J/J females and hair coat pigmentation was monitored in the progeny.

### Interspecies Chimeric Assays

Human iPSCs were cultured in serum-free N2B27-LCDM medium under 20% O_2_ and 5% CO_2_ at 37°C (Ref. ^7^). A total of 500 mL of N2B27 medium was prepared by including: 240 mL DMEM/F12 (Thermo Fisher Scientific, 11330-032), 240 mL Neurobasal (Thermo Fisher Scientific, 21103-049), 2.5 mL N2 supplement (Thermo Fisher Scientific, 17502-048), 5 mL B27 supplement (Thermo Fisher Scientific, 12587-010), 1% GlutaMAX (Thermo Fisher Scientific, 35050-061), 1% nonessential amino acids (Thermo Fisher Scientific, 11140-050), 0.1 mM β-mercaptoethanol (Thermo Fisher Scientific, 21985-023), and penicillin-streptomycin (Thermo Fisher Scientific, 15140-122). To prepare the N2B27-LCDM medium, small molecules and cytokines were added in the N2B27 medium as indicated at the following final concentrations: 10 ng/ml recombinant human LIF (L, Peprotech, 300-05), 1 μM CHIR99021 (C, Tocris, 4423), 2 μM (S)-(+)-Dimethindenemaleate (D, Tocris, 1425) and 2 μM Minocycline hydrochloride (M, Santa Cruz Biotechnology, sc-203339). Y-27632 (5 μM; Tocris, 1254) was added 24 h before and after single cell passage. Human iPSCs were cultured on mitomycin C (Sigma-Aldrich, M4287)-inactivated mouse embryonic fibroblast (MEF) feeder cells (3*10^4^ cells/cm^2^). The N2B27-LCDM medium was changed every day with fresh LCDM medium. To maintain human iPSCs in an undifferentiated state, we used the following criteria: a) avoid plating human cells too sparsely; b) use the proper quantity of freshly prepared MEF feeder cells; and c) do not allow human cells to overgrow. iPSCs were passaged by Accumax (Innovative Cell Technologies) every 3–5 days (normally at a split ratio ranging from 1:3 to 1:10), and the exact passage day and split ratio should be determined for each cell line specifically.

For transient miR-203 expression, human iPSCs expressing the Tomato reporter were transfected with pMCSV-miR203-GFP. The cell pellet was resuspended with the Nucleofection/Plasmid mixture (P3 Primary Cell 4D-Nucleofector Kit, Lonza): 82 μL solution + 18 μL supplement1 + 2 μg plasmid. The cells were transferred into a cuvette for nucleofection using a 4D-Nucleofector (Program CB150, Lonza), and then seeded on DR4 MEFs. The next day, the cells were gently digested into single cells using Accumax. Suspensions were filtered through a cell strainer (40 μm). Then, the samples were loaded and sorted to harvest GFP+ and Tomato+ cells on a Becton-Dickinson Fusion sorter. After cell collection, they were plated on DR4 MEFs in N2B27-LCDM medium plus10 μM Y-27632. The medium was changed the next day without Y-27632.

For interspecies chimeric experiments, human cells were used one day before passaging, which showed an optimal undifferentiated morphology and proliferated exponentially. At this time point, the colonies were at sub-confluent density (approximately 70% density of the day the cells should be passaged). 8-10-week ICR female mice were superovulated by the treatment of PMSG and 48 h later intraperitoneal administration of hCG. Female mice were then mated with a male for oocyte fertilization. Appearance of plugs were checked in female mice, and 2-cell stage embryos were harvested 1 day after that. 2-cell stage embryos were cultured in KSOM for another 24h and named E2.5 embryos. At this time most of the embryos were around 8-cell stage and ready for human EPS cell injection.

Human iPSCs were first rinsed with PBS, then treated with Accumax for 2 min and subsequently filtered through a cell strainer (40 μm). Cells were then centrifuged at 1,200-1,500 rpm (250-300 *g*) at room temperature for 3 min. The supernatant was removed, and the cells were re-suspended in the culture medium at a proper density (2-6*10^5^ cells/ml). 10 μM Y-27632 was added into the suspension. The suspension was placed on ice before injection. One individual cell was microinjected into each E2.5 ICR diploid mouse embryo. Injected embryos were cultured in N2B27-LCDM medium for the first 4 h (in the presence of 10 μM Y-27632), followed by culture for 44-60 h in mixed cell culture medium and KSOM medium at a 1:1 ratio. When the embryo reached the blastocyst stage, approximately 10 injected embryos were transferred to each uterine horn of a 2.5 days post-coitum pseudo-pregnant ICR female, or fixed with 4% paraformaldehyde for immunostaining. Postimplantation embryos were collected at E9.5, rinsed in PBS, followed by embedding, freezing, slicing (10 μm thick) with a Cryostat.

For immunofluorescence, embryos or slides were fixed in 4% paraformaldehyde (Sigma) at room temperature for 20 min and washed with PBST for 30 min. Samples were then permeabilized with 0.2% Triton X-100 in PBS. After 3 washings, samples were blocked with PBS (Corning, 21-040-CVR) that contained 1% BSA and 0.05% TWEEN-20 at room temperature for 45 min, and then incubated with primary antibodies at 4°C overnight. After 3 washings, secondary antibodies (Jackson ImmunoResearch) were incubated at room temperature for 1 hr. The nuclei were stained with DAPI (Southern Biotech). The following antibodies were used: anti-OCT4 (C30A3) (1:200; Cell Signaling), Anti-HuNu (1:200; Novus, NBP2-34342), anti-Gata4 (1:200; Santa Cruz, C-20), anti-Sox2 (1:200; R&D, AF2108), anti-hSOX17 (1:200; R&D, AF1924) (Supplementary Table 8). Immunofluorescence images were obtained using a confocal microscope and signals were then analyzed and quantified using ImageJ.

All the chimeric experiments were reviewed and approved by the Salk Institutional Animal Care and Use Committee (IACUC) and followed the ethical guidelines of the Salk Institute.

### Cell culture and gene expression

Primary mouse embryonic fibroblasts (MEFs) were obtained from embryos at E13.5 and cultured in DMEM supplemented with 10% of FBS and Penicillin/Streptomycin. Cultures were routinely tested for mycoplasma. Regarding the miR-203 inducible model-derived MEFs, reprogramming was promoted on these MEF cultures by non-inducible Oct4-Sox2-Klf4-cMyc (OSKM) lentiviral transduction. For lentiviral transduction, we transfected HEK293T cells with FUW-OSKM (Addgene #20328) and packaging vectors using Lipofectamine 2000 (Invitrogen). Viral supernatants were collected twice a day on two consecutive days starting 24 h after transfection and were used to infect MEFs, previously plated at a density of 250.000 cells per well in 6-well plates. Previous to infection, polybrene was added to the viral supernatants at a concentration of 2 μg/ml. For all those experiments in which miR-203 was expressed in WT iPSCs by mimics transfection or retrovirus transduction, WT MEFs reprogramming to obtain WT iPSCs was performed using inducible OSKM lentiviral transduction (TetO-FUW-OSKM; Addgene #20321). Infected MEFs were then cultured in iPSC medium, containing KO-DMEM (Gibco), 2-Mercaptoethanol 50 mM (Invitrogen), non-essential aminoacids MEM NEAA (Invitrogen), Penicillin and Streptomycin (5000 μg/ml, Invitrogen), Lif (Leukemia Inhibitor Factor, ESGRO, Millipore) and 20% knock-out serum replacement (KSR, Invitrogen). In this case, doxycycline was also added to the iPSC medium to promote reprograming only for the generation of WT iPSCs to be treated with miR-203 mimics or retroviruses (but never in the case of iPSCs derived from the miR-203 inducible animal model, in which miR-203 expression depends on DOX addition). Medium was changed every 24 h and plates were stained for alkaline phosphatase activity to assure the efficiency of reprogramming (AP detection kit, Sigma-Aldrich). Once colonies were picked, iPSCs were cultured in iPSCs media over mitomycin C (Roche)-inactivated feeder cells. G4 ESCs were cultured over mitomycin C-inactivated feeders and in the presence of ESC medium containing KO-DMEM, 2-Mercaptoethanol, non-essential amino acids, Glutamax, Penicillin and Streptomycin, Lif and 10% Fetal Calf Serum (Hyclone). When indicated, the culture media for pluripotent cells included 2i factors (MEK inhibitor PD0325901 1 μM, Gsk3 inhibitor CHIR99021 3 μM) and mouse LIF (as above) in N2B27 medium, as described previously^3^. For inducing transient miR-203 over-expression, *ColA1*(miR-203/miR-203); *Rosa26*(rtTA/rtTA) iPSC or ESC cultures were treated with Dox (1 μg/ml; Invitrogen) during 5 days. After that, Dox withdrawal was standardized for the cultures during following several passages (15-30 days) unless other time points are indicated in the text. In this inducible system, we always test that insert expression is uniquely dependent on Dox, and becomes absolutely undetectable after Dox withdrawal. As a control of the treatment itself, Dox was also added and tested in wild-type iPSCs or ESCs.

For over-expression experiments, miR-203 and full-length cDNAs of Dnmt3a and Dnmt3b were subcloned into the retroviral vector pMCSV-PIG^18^ by restriction-directed subcloning, using pCMV-Sport6-mDnmt3a (MGC clone:5662) and attB-mCh-mDnmt3b-Poly (A)-NeoR (Addgene plasmid 65553) as templates, respectively. For retroviral transduction, we transfected HEK293T cells with the respective pMCSV-PIG vector expressing a GFP reporter (including either only GFP; miR-203-GFP; Dnmt3a-GFP or Dnmt3b-GFP) and the packaging vector pCL-ECO, using Lipofectamine 2000 (Invitrogen). Viral supernatants were collected twice a day on two consecutive days starting 24 h after transfection and were used to infect either ESCs or IPSCs, previously plated over feeders in 6-well plates. Preceding the infection, polybrene was added to the viral supernatants at a concentration of 2 μg/ml. When transduced with these retroviral vectors, both ESCs or iPSCs cells were sorted by FACS and GFP-positive cells were selected for subsequent cultures and analysis. For transfection with mimics, we used miRIDIAN microRNA human hsa-miR-203a-5p (C-302893-00)/has-miR-203a-3p (C-300562-03) mimics or mimic transfection control (scramble sequences purchased from GE Dharmacon) with Dy547 (CP-004500-01), all of them from GE Dharmacon. Transfection was performed using either Dharmafect transfection reagent (Dharmacon) or Lipofectamine RNAiMAX (Invitrogen) following the manufacture’s instructions. Transfection efficiency was evaluated 24 h post-transfection by Dy547 fluorescence, and experiments were then performed as indicated in the figures.

For RNA interference assays, ON-TARGETplus SMARTpool for Non-targeting control siRNA (D-001810-01, 02, 03, 04), *Dnmt3a* siRNAs (J-065433-09, 10, 11, 12) and *Dnmt3b* siRNAs (J-044164-05, 06, 07, 08) from Dharmacon were used. Transfection was performed using Dharmafect transfection reagent (Dharmacon) following the manufacture’s instructions. Transfection efficiency was evaluated 24 h post-transfection by qPCR, using the primers indicated in Supplementary Table 9. A few weeks after transfection, the cells were assessed for differentiation to embryoid bodies.

Luciferase reported assays were performed in HEK293T cells. Briefly, 200.000 cells per well were seeded on 6-well plates and the day after, cells were transfected using Lipofectamine 2000 (Invitrogen), following the manufacture’s instructions. The 3’UTR regions from the murine genes *Dnmt3a, Dnmt3b, Dnmt3l* and *Dnmt1* were amplified by PCR with specific primers (Dnmt3a_EcoRI-Fw: 5’-GAATTCAGGGACATGGGGGCAAACTGAA-3’ (SEQ ID NO:55); Dnmt3a_NdeI-Rv: 5’-CATATGCTGAGGCAGTCATTTTAGATTCAT-3’ (SEQ ID NO:56); Dnmt3b_EcoRI-Fw: 5’-GAATTCTTTTAGCTCACCTGTGTGGGG-3’ (SEQ ID NO:57); Dnmt3b_NdeI-Rv: 5’-CATATGCCAGAAAGGTAAACTCTGGGCA-3’ (SEQ ID NO:58); Dnmt3l_EcoRI-Fw: 5’-GAATTCGAAATGAATCACCATAAGATGAAAG-3’(SEQ ID NO:59); Dnmt3l_NdeI-Rv: 5’-CATATGAACAATCCTATGATATATTTGAAAAA-3’ (SEQ ID NO:60); Dnmt1_EcoRI-Fw: 5’-GAATTCGTGCTCTCACCCAGAGCCCCA-3’ (SEQ ID NO:61); Dnmt1_NdeI-Rv: 5’-CATATGGCTTGACAGAAGCGCTTTATTTTG-3’) (SEQ ID NO:62)(Supplementary Table 9), using cDNA clones (pCMV-Sport6-mDnmt3a, cDNA clone MGC:5662; pBluescript-mDnmt3l, RIKEN clone: 2410006021; pYX-Asc-mDnmt1, MGC clone:62302) or mouse cDNA (in the case of *Dnmt3b*). PCR products were verified by sequencing and ligated into the pGL3-Control vector (Promega), downstream of the luciferase reporter gene. Mutations in the miR-203 binding sites (Supplementary Fig. 9) were generated by site-directed mutagenesis and subsequently verified by sequencing. Transfections were performed with either pMCSV-GFP or pMCSV-miR-203-GFP vectors, in combination with the pGL3-derived vectors, and Renilla as a control. Luciferase measurement was achieved 48 h post-transfection using a luminescence microplate reader (Biotek). Experimentation with human cells was performed according to protocols approved by the ISCIII Ethics Committee for Research (CEI; number PI 61_2017).

### Embryoid body generation

Briefly, iPSCs or ESCs were trypsinized and resuspended to a concentration of 200,000 cells/ml in the presence of complete growth medium lacking leukemia inhibitory factor (Lif). Small drops of this suspension (∼35 μl) were collected and seeded on the lid of a 10 mm plate, generating hanging drops of approximately 5,000 cells per drop. After 4 days, the aggregates were already visible at the bottom of the drops and were picked and transferred to a non-adherent plate, containing DMEM and 10% Fetal Calf Serum. There they were maintained in suspension for the indicated times and beating, size and cavity formation were assessed every five days as described in the figures. Between 20 and 30 EBs were analyzed per condition and the percentage of beating EBs or EBs with large cavities were calculated for every time point. EB size was measured using the Image J software.

### Immunofluorescence and immunohistochemistry

Cells previously seeded in cover slips were fixed in 4% paraformaldehyde for 15 min, permeabilized using PBS 0.1% Triton X-100 for 15 min and blocked in BSA for 1 h at room temperature. Primary antibody incubation was performed overnight at 4°C in all cases, followed by secondary antibody incubation for 1 hour at room temperature. Nuclear staining was included in the last PBS wash, using Hoescht or DAPI. Primary antibodies used in this study were against Cd34 (Abcam), Gata4 (Santa Cruz), cTnT (Abcam) and phospho-histone H3 (Millipore) (Supplementary Table 8). Cells were examined under a Leica SP5 microscope equipped with white light laser and hybrid detection.

For immunohistochemistry, tissue samples were fixed in 10% neutral buffered formalin (4% formaldehyde in solution), paraffin-embedded and cut at 3 µm, mounted in superfrost®plus slides and dried overnight. Consecutive sections were stained with hematoxylin and eosin (H&E) or subjected to immunohistochemistry using automated immunostaining platforms (Ventana Discovery XT, Roche or Autostainer Plus Link 48). Antigen retrieval was first performed with high or low pH buffer depending on the primary antibody (CC1m, Roche or low pH antigen retrieval buffer, Dako), endogenous peroxidase was blocked (peroxide hydrogen at 3%) and slides were incubated with primary antibodies against cytokeratin 8 (CK8; CNIO Monoclonal Antibodies Core Unit, AM-TROMA I), GFP (Roche, 11814460001), Sox2 (Cell Signaling Technology, 3728), alpha-fetoprotein (AFP; R&D Systems, AF5369), Oct4 (Santa Cruz Biotechnology, sc-9081), Cd34 (ABCAM ab8158), Gata4 (Santa Cruz Biotechnology, sc-1237), or Ter119 (LY-76; BD Bioscience, 550565) (Supplementary Table 8). Secondary antibodies were conjugated with horseradish peroxidase (OmniRabbit, Ventana, Roche) and the immunohistochemical reaction was developed using 3,30-diaminobenzidine tetrahydrochloride (DAB) as a chromogen (Chromomap DAB, Ventana, Roche or DAB solution, Dako) and nuclei were counterstained with Carazzi’s hematoxylin. Finally, the slides were dehydrated, cleared and mounted with a permanent mounting medium for microscopic evaluation. For Sirius red staining, Weigert’s Hematoxylin was incubated for 8 min and Picro/Sirius Red for 1 h, followed by a wash (10 min) with water. The images were acquired with a slide scanner (AxioScan Z1, Zeiss). Sirius red staining was measured using both bright-field and polarized lights. Images were captured and quantified using the Zen Software (Zeiss).

### Analysis of mRNA and microRNA levels

RNA was extracted from cell or tissue samples with Trizol (Invitrogen) or by using the miRVana miRNA isolation kit (Thermo Fisher), following the manufacture’s recommendations. Retrotranscription into cDNA was performed using M-MLV Reverse Transcriptase (Promega) following the manufacturer’s protocol. Quantitative real time PCR was performed using Syber Green Power PCR Master Mix (Applied Biosystems) in an ABI PRISM 7700 Thermocycler (Applied Biosystems). The housekeeping gene *Gapdh* was used for normalization. The oligonucleotide primers used in this manuscript are listed in the Supplementary Table 9. For reverse transcription of microRNAs, we used the Taqman small RNA assay (4366596), including the specific oligonucleotides for mmu-miR-203-5p and 3p (002580 and 000507), miR-16 and the housekeeping RNAs sno-202 or sno-142. Conditions for miRNA amplification were as follows: 30 minutes at 16°C; 30 minutes at 42°C and a final step of 5 minutes at 85°C. Quantitative real time PCR was then performed using the Taqman Universal PCR Master Mix (434437) following the manufacture’s instructions in an ABI PRISM 7700 Thermocycler (Applied Biosystems).

For RNAseq, total RNA was extracted using the miRVana miRNA isolation kit (ThermoFisher), following the manufacture’s recommendations. Between 0.8 and 1 µg of total RNA was extracted from iPSCs, ESCs or teratomas, with RIN (RNA integrity number) numbers in the range of 9 to 10 (Agilent 2100 Bioanalyzer). Poly A+ fractions were purified and randomly fragmented, converted to double stranded cDNA and processed using the Illumina’s “TruSeq Stranded mRNA Sample Preparation Part # 15031047 Rev. D” kit. The adapter-ligated library was completed by PCR with Illumina PE primers (8-11 cycles) and the resulting directional cDNA libraries were sequenced for 50 bases in a single-read format (Illumina HiSeq2000) and analyzed with nextpresso ^50^. Reads were quality-checked with FastQC (http://www.bioinformatics.babraham.ac.uk/projects/fastqc) and aligned to the mouse genome (GRCm38/mm10) with TopHat-2.0.10 ^51^, using Bowtie 1.0.0 ^52^ and Samtools 0.1.19 ^53^, allowing two mismatches and five multihits. Transcripts assembly, estimation of their abundances and differential expression were calculated with Cufflinks 2.2.1 ^51^, using the mouse genome annotation data set GRCm38/mm10 from the UCSC Genome Browser^54^. A false discovery rate (FDR) of 0.05 is used as threshold for significance in differential expression. Heatmaps were later built using GENE-E (http://www.broadinstitute.org/cancer/software/GENE-E/index.html).

### Bisulphite conversion, genome-wide DNA methylation and validation of DMRs

DNA samples were prepared for whole genome bisulphite sequencing using the TrueMethyl® Whole Genome Kit (CEGX®) according to the manufacturer’s instructions. Briefly, 200 ng of genomic DNA was sheared to 800 bp using M220 Focused-ultrasonicator™(Covaris®). Then the fragmented DNA was denatured and oxidised by a chemical oxidant to convert 5-hydroxymethyl cytosine to 5-formylcytosine (5fC). The purpose of the oxidation was to ensure that we purely captured the information for 5’ methyl cytosine methylation and not an indistinguishable pattern combination of 5’methyl cytosine and 5’hydroxymethyl cytosine. Following oxidation, the DNA was subjected to bisulphite conversion for the deamination of cytosines and 5fC to uracils. Bisulphite converted DNA was desulfonated and purified to then proceed to library preparation. In this “post-bisulphite conversion” library preparation method, the fragmented single stranded bisulphite converted DNA was adapted with sequencing adaptors at the 3’end followed by an extension step and finally ligation of adaptors at the 5’ end of the molecules. Finally, the libraries were indexed and amplified. The PCR was performed for 10 cycles followed by bead-based purification. An additional purification and size selection step using Agencourt AMPure XP beads (Beckman Coulter, Cat: A63881) was performed to remove adaptor dimers. The purified library was eluted in a final volume of 14 μl pure water. Quality of the library obtained was checked using the DNA High sensitivity chip on the Agilent Bioanalyzer. The library was quantified using Qubit and KAPA Biosystems Library quantification kit (#KK4824) according to manufacturer’s instructions. A total of 1-2pMol of multiplexed libraries were sequenced on a 125 bp paired-end format (Illumina HiSeq2500). Adapter sequences were removed using cutadapt version 1.9.1^55^ in paired-end mode with parameters “-m 15 -u 6 -U 6”. Bwa-meth^56^ was then used to align reads to mm10 using default parameters. PCR duplicates were removed using Picard v1.91 (http://broadinstitute.github.io/picard). Count tables of the number of methylated and unmethylated bases sequenced at each CpG site in the genome were constructed using the “tabulate” module of bwa-meth and BisSNP-0.82.2 ^57^ with default parameters. All libraries passed basic quality control checks with a minimum of 82.7% of read pairs aligning uniquely. DMRs were called using the WGBS module of DMRcate ^58^ with parameters lambda=1000 and C=50, DMPs were called using DSS^59^, PMDs, LMRs and UMRs were called using MethylSeekR^60^ and PMDs were called using the R package ‘aaRon’ (https://github.com/astatham/aaRon).

For the validation of DMRs, clonal bisulphite sequencing was performed at two particular loci: *Elf5* (chr2: 103, 423, 778-103, 424, 180) and *Sirt6* (chr10: 81, 624, 595-81, 625, 547) promoter regions. 100 ng of DNA was bisulphite treated using EZ DNA-Methylation-Lighting^TM^ kit (Zymo Research) following manufacturer’s instructions. Triplicate PCR amplifications were then performed using semi-nested bisulphite conversion specific primers listed in Supplementary Table 9, following the PCR conditions previously described for *Elf5* ^61^ or using the following protocol: 95°C 4 min; 5 cycles (95°C 45 sec, 54°C 1.5 min, 72°C 2 min); 25 cycles (95°C 45 sec, 54°C 1.5 min; 72°C 1.5 min) and final extension at 72°C 4 min, for *Sirt6*. The methylation status of the PCR amplicons was determined by Sanger clonal sequencing of the pooled PCR products to ensure representative clonal analysis using between 8-10 clones per sample. The analysis of the methylation status of the clones was performed using BiQ Analyzer^62^.

### Neonatal cardiomyocyte isolation and differentiation studies

Neonatal mouse and rat cardiomyocytes were prepared as described previously^63^. Briefly, neonatal cardiomyocytes were isolated by enzymatic disassociation of 1-day-old neonatal mouse or rat hearts with the Neonatal Cardiomyocyte Isolation System (Cellutron Life Technology). Cardiomyocytes were pre-plated for 2 hours to remove fibroblasts. Cells were then plated on 1% gelatin-coated plates in medium containing 10% horse serum and 5% fetal calf serum. Eighteen hours after plating, cells were changed into serum-free medium and then transfected with 50 nM miR-203a-3p & 5p mimics or control mimics (from Dharmacon) by using Lipofectamine RNAiMAX transfection reagent, following the manufacture’s instructions. Six hours later, media with transfection reagent were removed and substituted by media containing 1% serum. Different time points (as indicated) were collected for subsequent RNA extraction and EdU staining/immunofluorescence. For EdU staining, we used the Click-iT EdU staining Kit (Invitrogen) following the manufacture’s recommendations. After that, immunofluorescence of cardiomyocytes markers (Troponin T) was performed and Hoescht was finally used for nuclei staining.

For in vitro differentiation from mouse iPSCs to cardiomyocytes, wild-type mouse iPSCs were transfected with either control or miR-203 mimics prior to differentiation. In some experiments, additionally pMCSV-Dnmt3a and 3b or empty pMCSV vector were transiently transduced, 24 hours after mimics transfection. iPSCs were then maintained in culture under 2i/Lif media for 15 days, before differentiation started. Then, cells were plated in Matrigel pre-coated 6-well plates. Once they become confluent, the media was changed to RPMI 1640/B27 medium (Thermo Fisher Scientific, 61870127 for the RPMI 1640 media and A1895601 for B27) with CHIR99021 5 μM (Stem Cell Technologies, 72054). After two days, cells were washed and media without CHIR99021 was added. At day 3 of differentiation, media was supplemented with IWR1 5 μM (Stem Cell Technologies, 72564) and maintained in these conditions for 2 more days. At day 7 of differentiation, the media was changed to RPMI 1640/B27 plus insulin. Then, from day 9 to day 15, cells were cultured in RPMI 1640/B27 and the media was changed every 2 days. From day 16 to day 18 of differentiation, cells were cultured in DMEM without glucose and with lactate 4mM. From day 18 to day 21, the media was changed to RPMI 1640/B27 and changed every day. Finally, from day 22 to day 28, monolayer cells can be isolated into cell clusters and kept in low attachment for another week.

### Cardiac cryoinjury in neonatal mice

The neonatal mouse cardiac cryoinjury experiments were performed as described previously^34^. Briefly, at postnatal day 1, neonates were placed on a heated water blanket set to 37 °C and covered with bedding from mothe’s nest. They were then anesthetized by placing the pups into an ice-water bath for 3 minutes. After that, the pups were dried using a sterile gauze pad and placed in the surgical area in the supine position, immobilized at the arms legs and tail. A transverse skin incision across the chest was made using a pair of micro-scissors and then carefully, a lateral thoracotomy was performed, by making a small incision at the fourth or fifth intercostal space. The pericardial sac was then carefully removed and the heart was exteriorized by gently pressing the abdomen. The left ventricle was identified and then the precooled cryoprobe was applied on the ventricle surface for just 2 seconds. To close the chest wall we used 8-0 Non-absorbable Prolene sutures and to close the skin, we used Webglue sutures. After closure, the surgical area was washed with wet gauzes to remove any residual blood. Rapidly, the pup was warmed for several minutes, returned to the heating blanket with the other pups and covered with bedding from the mothe’s cage. Once all the pups had fully recovered from the surgery, they were swapped in the mothe’s cage. Seven days after the cryoinjury, the pups were euthanized by decapitation and the hearts were collected and fixed in formaldehyde. The hearts were then processed normally for paraffin embedding as indicated above.

### Statistics

Normal distribution and equal variance was confirmed before using the Student’s t-test (two-tailed, unpaired) to estimate statistical significance. For contingency tables, we used the Fisher’s exact test. Statistical analysis was performed using Prism (GraphPad Software, La Jolla, CA).

## Data availability

RNAseq and methylation data has been deposited in the GEO repository under accession numbers GEO SuperSeries GSE81571 and GSE86653.

## ACKNOWLEDGEMENTS

We thank María Guirola, Sheila Rueda and the CNIO Histopathology Unit for technical support, Carmen Gómez and Patricia Prieto for help with iPSC/ESC culture, Zsuzsanna Izsvak (Max-Delbrück-Center for Molecular Medicine, Berlin, Germany) for reagents, Marta Cañamero (Pathology and Tissue Analytics, Roche Pharma, Basel, Switzerland) and Alba de Martino (Histopathology Unit, CNIO) for pathology advice; and William Pu, Zhang-Peng Huang and Kai Li (Cardiovascular Research Division, Boston Childre’s Hospital, USA) for help with the cardiac differentiation experiments. We are indebted to the members of the CNIO Cell Division and Cancer laboratory for their constant support and advice. M.S.R. was supported by the Asociación Española contra el Cáncer (AECC; 2012 AIOA120833SALA and 2018 INVES18005SALA) and a Juan de la Cierva contract from the Spanish Ministry of Science (MICINN). M.T. was supported by Fundación La Caixa. M.A.F. was supported by MICINN (SAF2014-60442-JIN, cofunded by ERDF-EU. J.F.P. was funded by MICINN (RTI2018-093330-B/FEDER-EU), Ramón Areces Foundation (CIVP19S7917), CAM (B2017/BMD-3778) and AECC (2018, PROYE18054PIRI). F.V.M was supported by the National Breast Cancer Foundation/Cure Cancer Australia Foundation Postdoctoral Training Fellow. Work in S.J.C laboratory was supported by the National Health and Medical Research Council (NHMRC 1063560). M.A.B. laboratory is funded by SAF2013-45111-R from MICINN, Fundación Botín and Banco Santander, and Worldwide Cancer Research (WCR16-1177). Work in S.O. laboratory was funded by MICINN grant (SAF2013-44866-R). Work in the M.M. laboratory was supported by grants from the MICINN (SAF2015-69920-R cofunded by ERDF-EU), the iLUNG Programme (B2017/BMD-3884) from the Comunidad de Madrid, and the MicroKin network (ERA-NET NEURON & PCIN-2015-007). The CNIO is a Severo Ochoa Centers of Excellence (MINECO award SEV-2015-0510).

## AUTHOR CONTRIBUTION

M.S.R. performed the cellular and in vivo assays. M.T. contributed to the characterization of *mi*PSCs and M.A.F. and E.Z. to the generation of DNA constructs and molecular biology studies. F.V.M., T.J.P. and S.J.C. performed and analyzed the genome-wide methylation studies. C.Z., Y.Y. and J.C.I.B. performed the human-mouse interspecies chimeric assays. O.G. analyzed RNAseq data. M.A. and M.S. participated in the evaluation of the data. M.J.B. and M.G.C. generated the miR-203 knockin mouse model, and J.F.P. contributed to its analysis. R.S. and M.A.B. contributed to the cardiac cryoinjury experiments. D.Z.W. contributed to the analysis of cardiomyocyte differentiation. J.M. and S.O. performed and analyzed chimera and tetraploid complementation assays and S.O. supervised and analyzed embryo work. M.M. conceived the project and M.S.R. and M.M. supervised the work and wrote the manuscript with the help of the rest of the authors.

## REFERENCES

1. Takahashi, K. & Yamanaka, S. A decade of transcription factor-mediated reprogramming to pluripotency. Nature reviews. Molecular cell biology 17, 183–193 (2016).

2. Li, M. & Izpisua Belmonte, J.C. Looking to the future following 10 years of induced pluripotent stem cell technologies. Nature protocols 11, 1579–1585 (2016).

3. Ying, Q.L. et al. The ground state of embryonic stem cell self-renewal. Nature 453, 519–523 (2008).

4. Choi, J. et al. Prolonged Mek1/2 suppression impairs the developmental potential of embryonic stem cells. Nature 548, 219–223 (2017).

5. Yagi, M. et al. Derivation of ground-state female ES cells maintaining gamete-derived DNA methylation. Nature 548, 224–227 (2017).

6. Choi, Y.J. et al. Deficiency of microRNA miR-34a expands cell fate potential in pluripotent stem cells. Science 355 (2017).

7. Yang, Y. et al. Derivation of Pluripotent Stem Cells with In Vivo Embryonic and Extraembryonic Potency. Cell 169, 243–257 e225 (2017).

8. Yi, R., Poy, M.N., Stoffel, M. & Fuchs, E. A skin microRNA promotes differentiation by repressing ‘stemness’. Nature 452, 225–229 (2008).

9. Bueno, M.J. et al. Genetic and epigenetic silencing of microRNA-203 enhances ABL1 and BCR-ABL1 oncogene expression. Cancer cell 13, 496–506 (2008).

10. Michel, C.I. & Malumbres, M. microRNA-203: Tumor Suppression and Beyond. MicroRNA 2, 118–126 (2013).

11. Yang, Y. et al. Determination of microRNAs in mouse preimplantation embryos by microarray. Developmental dynamics : an official publication of the American Association of Anatomists 237, 2315–2327 (2008).

12. Goossens, K. et al. Regulatory microRNA network identification in bovine blastocyst development. Stem cells and development 22, 1907–1920 (2013).

13. Takahashi, K. & Yamanaka, S. Induction of pluripotent stem cells from mouse embryonic and adult fibroblast cultures by defined factors. Cell 126, 663–676 (2006).

14. Stadtfeld, M., Maherali, N., Borkent, M. & Hochedlinger, K. A reprogrammable mouse strain from gene-targeted embryonic stem cells. Nature methods 7, 53–55 (2010).

15. Chung, H.C. et al. Human induced pluripotent stem cells derived under feeder-free conditions display unique cell cycle and DNA replication gene profiles. Stem cells and development 21, 206–216 (2012).

16. Biase, F.H., Cao, X. & Zhong, S. Cell fate inclination within 2-cell and 4-cell mouse embryos revealed by single-cell RNA sequencing. Genome research 24, 1787–1796 (2014).

17. Eckersley-Maslin, M. et al. Dppa2 and Dppa4 directly regulate the Dux-driven zygotic transcriptional program. Genes & development 33, 194–208 (2019).

18. Abad, M. et al. Reprogramming in vivo produces teratomas and iPS cells with totipotency features. Nature 502, 340–345 (2013).

19. Li, Y. et al. Generation of iPSCs from mouse fibroblasts with a single gene, Oct4, and small molecules. Cell Res 21, 196–204 (2011).

20. Sandhu, R., Rivenbark, A.G. & Coleman, W.B. Loss of post-transcriptional regulation of DNMT3b by microRNAs: a possible molecular mechanism for the hypermethylation defect observed in a subset of breast cancer cell lines. International journal of oncology 41, 721–732 (2012).

21. Gasque Schoof, C.R., Izzotti, A., Jasiulionis, M.G. & Vasques Ldos, R. The Roles of miR-26, miR-29, and miR-203 in the Silencing of the Epigenetic Machinery during Melanocyte Transformation. BioMed research international 2015, 634749 (2015).

22. To, K.K., Leung, W.W. & Ng, S.S. A novel miR-203-DNMT3b-ABCG2 regulatory pathway predisposing colorectal cancer development. Molecular carcinogenesis 56, 464–477 (2017).

23. Xie, W. et al. Epigenomic analysis of multilineage differentiation of human embryonic stem cells. Cell 153, 1134–1148 (2013).

24. Lister, R. et al. Human DNA methylomes at base resolution show widespread epigenomic differences. Nature 462, 315–322 (2009).

25. Latos, P.A. et al. Elf5-centered transcription factor hub controls trophoblast stem cell self-renewal and differentiation through stoichiometry-sensitive shifts in target gene networks. Genes & development 29, 2435–2448 (2015).

26. Chen, T., Ueda, Y., Dodge, J.E., Wang, Z. & Li, E. Establishment and maintenance of genomic methylation patterns in mouse embryonic stem cells by Dnmt3a and Dnmt3b. Molecular and cellular biology 23, 5594–5605 (2003).

27. Blattler, A. et al. Global loss of DNA methylation uncovers intronic enhancers in genes showing expression changes. Genome biology 15, 469 (2014).

28. Liao, J. et al. Targeted disruption of DNMT1, DNMT3A and DNMT3B in human embryonic stem cells. Nature genetics 47, 469–478 (2015).

29. Mikkelsen, T.S. et al. Dissecting direct reprogramming through integrative genomic analysis. Nature 454, 49–55 (2008).

30. Jackson, M. et al. Severe global DNA hypomethylation blocks differentiation and induces histone hyperacetylation in embryonic stem cells. Molecular and cellular biology 24, 8862–8871 (2004).

31. Miyata, S., Minobe, W., Bristow, M.R. & Leinwand, L.A. Myosin heavy chain isoform expression in the failing and nonfailing human heart. Circulation research 86, 386–390 (2000).

32. Kattman, S.J. et al. Stage-specific optimization of activin/nodal and BMP signaling promotes cardiac differentiation of mouse and human pluripotent stem cell lines. Cell stem cell 8, 228–240 (2011).

33. Otsuji, T.G. et al. Progressive maturation in contracting cardiomyocytes derived from human embryonic stem cells: Qualitative effects on electrophysiological responses to drugs. Stem cell research 4, 201–213 (2010).

34. Polizzotti, B.D., Ganapathy, B., Haubner, B.J., Penninger, J.M. & Kuhn, B. A cryoinjury model in neonatal mice for cardiac translational and regeneration research. Nature protocols 11, 542–552 (2016).

35. Zhao, X.Y. et al. iPS cells produce viable mice through tetraploid complementation. Nature 461, 86–90 (2009).

36. Kang, L., Wang, J., Zhang, Y., Kou, Z. & Gao, S. iPS cells can support full-term development of tetraploid blastocyst-complemented embryos. Cell stem cell 5, 135–138 (2009).

37. Mascetti, V.L. & Pedersen, R.A. Human-Mouse Chimerism Validates Human Stem Cell Pluripotency. Cell stem cell 18, 67–72 (2016).

38. Wu, J., et al. Stem cells and interspecies chimaeras. Nature 540, 51–59 (2016).

39. Okita, K., Ichisaka, T. & Yamanaka, S. Generation of germline-competent induced pluripotent stem cells. Nature 448, 313–317 (2007).

40. Stadtfeld, M. et al. Aberrant silencing of imprinted genes on chromosome 12qF1 in mouse induced pluripotent stem cells. Nature 465, 175–181 (2010).

41. Leitch, H.G. et al. Naive pluripotency is associated with global DNA hypomethylation. Nat Struct Mol Biol 20, 311–316 (2013).

42. Theunissen, T.W. et al. Molecular Criteria for Defining the Naive Human Pluripotent State. Cell stem cell 19, 502–515 (2016).

43. Okano, M., Bell, D.W., Haber, D.A. & Li, E. DNA methyltransferases Dnmt3a and Dnmt3b are essential for de novo methylation and mammalian development. Cell 99, 247–257 (1999).

44. Hirasawa, R. et al. Maternal and zygotic Dnmt1 are necessary and sufficient for the maintenance of DNA methylation imprints during preimplantation development. Genes & development 22, 1607–1616 (2008).

45. Branco, M.R., Oda, M. & Reik, W. Safeguarding parental identity: Dnmt1 maintains imprints during epigenetic reprogramming in early embryogenesis. Genes & development 22, 1567–1571 (2008).

46. Polizzotti, B.D. et al. Neuregulin stimulation of cardiomyocyte regeneration in mice and human myocardium reveals a therapeutic window. Science translational medicine 7, 281ra245 (2015).

47. Bar-Nur, O. et al. Small molecules facilitate rapid and synchronous iPSC generation. Nature methods 11, 1170–1176 (2014).

48. Chen, J. et al. Vitamin C modulates TET1 function during somatic cell reprogramming. Nature genetics 45, 1504–1509 (2013).

49. Beard, C., Hochedlinger, K., Plath, K., Wutz, A. & Jaenisch, R. Efficient method to generate single-copy transgenic mice by site-specific integration in embryonic stem cells. Genesis 44, 23–28 (2006).

50. Graña, O., Rubio-Camarillo, M., Fernandez-Riverola, F., Pisano, S.G. & Gonzalez-Peña, D. nextpresso: next generation sequencing expression analysis pipeline. Curr Bioinformatics 12, in press (2017).

51. Trapnell, C. et al. Differential gene and transcript expression analysis of RNA-seq experiments with TopHat and Cufflinks. Nature protocols 7, 562–578 (2012).

52. Langmead, B., Trapnell, C., Pop, M. & Salzberg, S.L. Ultrafast and memory-efficient alignment of short DNA sequences to the human genome. Genome biology 10, R25 (2009).

53. Li, H. & Durbin, R. Fast and accurate short read alignment with Burrows-Wheeler transform. Bioinformatics 25, 1754–1760 (2009).

54. Rosenbloom, K.R. et al. The UCSC Genome Browser database: 2015 update. Nucleic acids research 43, D670–681 (2015).

55. Martin, M. Cutadapt removes adapter sequences from high-throughput sequencing reads. EMBnet J 17, 10–12 (2011).

56. Pedersen, B.S., Eyring, K., De, S., Yang, I.V. & Schwartz, D.A. Fast and accurate alignment of long bisulfite-seq reads. arXiv, 1401–1129 (2014).

57. Liu, Y., Siegmund, K.D., Laird, P.W. & Berman, B.P. Bis-SNP: combined DNA methylation and SNP calling for Bisulfite-seq data. Genome biology 13, R61 (2012).

58. Peters, T.J. et al. De novo identification of differentially methylated regions in the human genome. Epigenetics & chromatin 8, 6 (2015).

59. Feng, H., Conneely, K.N. & Wu, H. A Bayesian hierarchical model to detect differentially methylated loci from single nucleotide resolution sequencing data. Nucleic acids research 42, e69 (2014).

60. Burger, L., Gaidatzis, D., Schubeler, D. & Stadler, M.B. Identification of active regulatory regions from DNA methylation data. Nucleic acids research 41, e155 (2013).

61. Lee, H.J. et al. Lineage specific methylation of the Elf5 promoter in mammary epithelial cells. Stem cells 29, 1611–1619 (2011).

62. Bock, C. et al. BiQ Analyzer: visualization and quality control for DNA methylation data from bisulfite sequencing. Bioinformatics 21, 4067–4068 (2005).

63. Huang, Z.P. et al. Cardiomyocyte-enriched protein CIP protects against pathophysiological stresses and regulates cardiac homeostasis. The Journal of clinical investigation 125, 4122–4134 (2015).

