## Supplementary Information for "A novel microRNA-based strategy to expand the differentiation potency of stem cells"

By Salazar-Roa et al.

### List of Expanded Material

#### Expanded views

Figure EV1. Mouse alleles generated and used in this work, transcriptomic analysis of *mi*i*PSCs* and validation of miR-203 mimics and vectors on EBs differentiation.

Figure EV2. Histopahological analysis of teratomas generated from *mi*iPSCs.

Figure EV3. miR-203-exposed human iPSCs efficiently contribute to human-mouse interspecies chimeras.

Figure EV4. miR-203 targets *Dnmt3a/b* 3'-UTR sequences.

Figure EV5. Genome-wide methylation of iPSCs or embryoid bodies after transient exposure to miR-203.

Figure EV6. Improved cardiomyocyte differentiation and maturation after transient expression of miR-203.

#### Supplementary Tables

Supplementary Table 1. Gene Ontology Analysis of genes upregulated in *mi*iPSCs.

Supplementary Table 2. Gene Ontology Analysis of genes significantly deregulated in *mi*iPSC-derived teratomas vs. un-induced iPSC-derived teratomas.

Supplementary Table 3. List of miR-203 predicted targets among the transcripts downregulated in *mi*iPSCs and involved in the epigenetic regulation of gene expression (GO0040029).

Supplementary Table 4. Differentially methylated positions (DMP) in *mi*iPSC (t=10 versus t=0 and t=25 versus t=0)

Supplementary Table 5. Extended data of genome-wide methylation analysis.

Supplementary Table 6. List of the 282 genes included in the 2-cell-embryo signature and their TSS methylation level in *mi*iPSCs

Supplementary Table 7. A comparison between *mi*PSCs and other PSCs previously published in representative studies.

Supplementary Table 8. Antibodies used in this work.

Supplementary Table 9. Oligonucleotides used in this work.

### Supplementary Figures

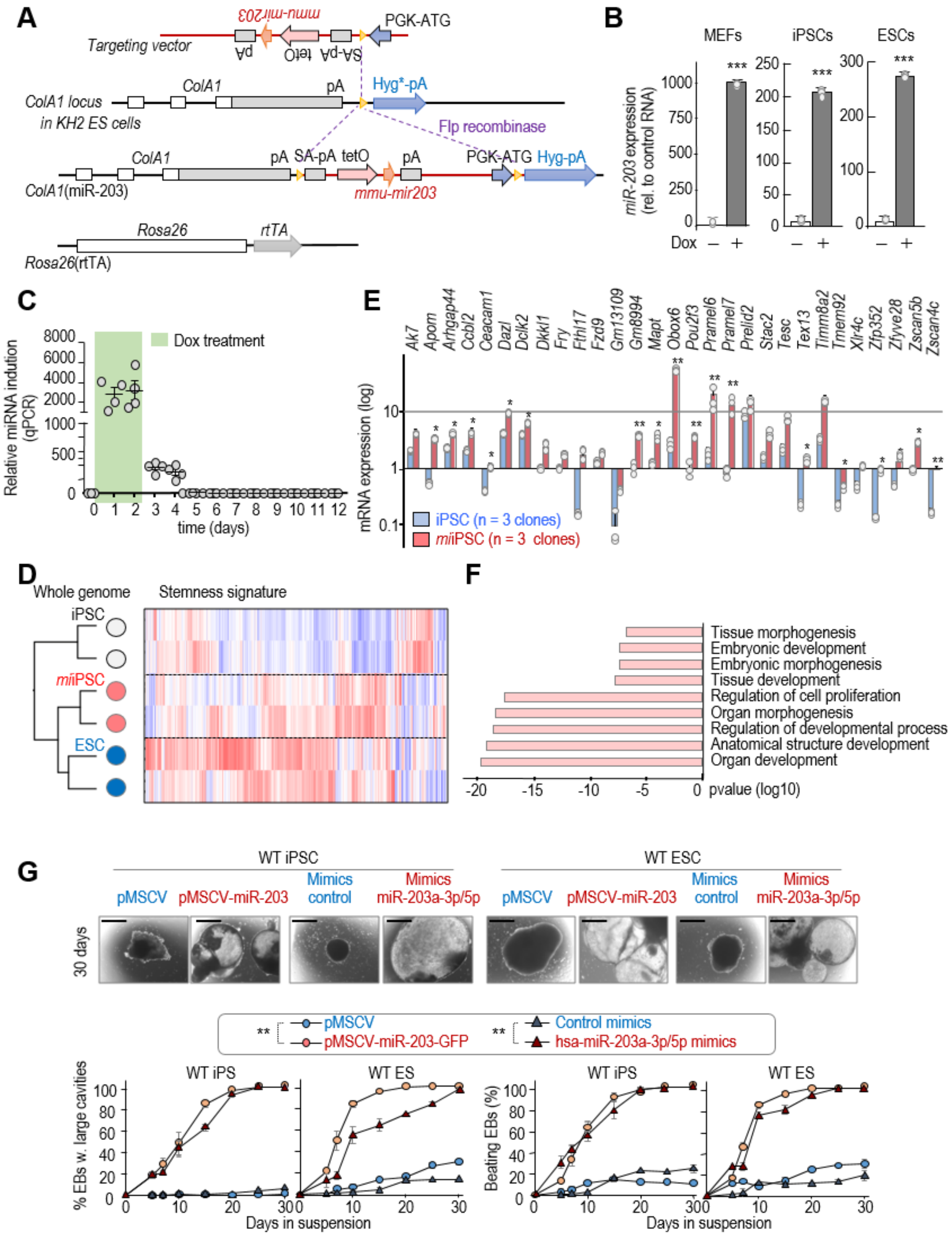

Figure EV1 Mouse alleles generated and used in this work, transcriptomic analysis of *mi*iPSCs and validation of miR-203 mimics and vectors on EBs differentiation. A, Schematic representation of the miR-203 inducible knockin model generated in this work. In the *ColA1*(miR-203/miR-203); *Rosa26*(rtTA/rtTA) model, the reverse tetracycline transactivator is expressed from the *Rosa26* locus, whereas miR-203 is driven by the tetracycline operator downstream of the *ColA1* locus. B, miR-203 expression, as determined by quantitative PCR, in *ColA1*(miR-203/miR-203); *Rosa26*(rtTA/rtTA) MEFs, iPSCs and ESCs treated or not with doxycycline (Dox). miR-203 expression is normalized by a control miRNA (miR-142). ***$P$<0.001

(Student´s *t*-test). C, Time course of miR-203 expression, as determined by qPCR. RNA expression is normalized by a housekeeping miRNA (miR-16) that maintained invariable. Data represent 4 different qPCR measures. *P*<0.001 (Student´s t-test) comparing Dox treatment (orange box) versus Dox withdrawal. D Unbiased clustering of genome-wide RNAseq data (left) and heatmap plot showing the comparative expression of 450 genes associated with pluripotency (Ref. [17]); 2 clones per sample. E, Early expression (RPKM) of the indicated transcripts included in the 2C-signature. Data are mean ± s.e.m (n=3 independent experiments). **P*<0.05; ***P*<0.01 (Student´s t-test). F, Top categories in the Gene Ontology Analysis of the genes significantly upregulated in *mi*iPSCs versus un-induced iPSCs (4 independent clones were analyzed; see also Supplementary Table 1). G, (Upper panels) Representative images of EBs derived from either wild-type iPSCs (left) or ESCs (right) transduced with empty pMCSV vector, pMCSV-miR-203 or transfected with control mimics or miR-203 mimics, at day 30 of the differentiation process. Scale bars, 500 µm. (Lower panels) Quantification of EBs with large cavities and beating EBs during the differentiation process. Data are mean ± s.e.m (n=3 independent experiments). ***P*<0.01 (both in iPS and ES cells; Student´s t-test).

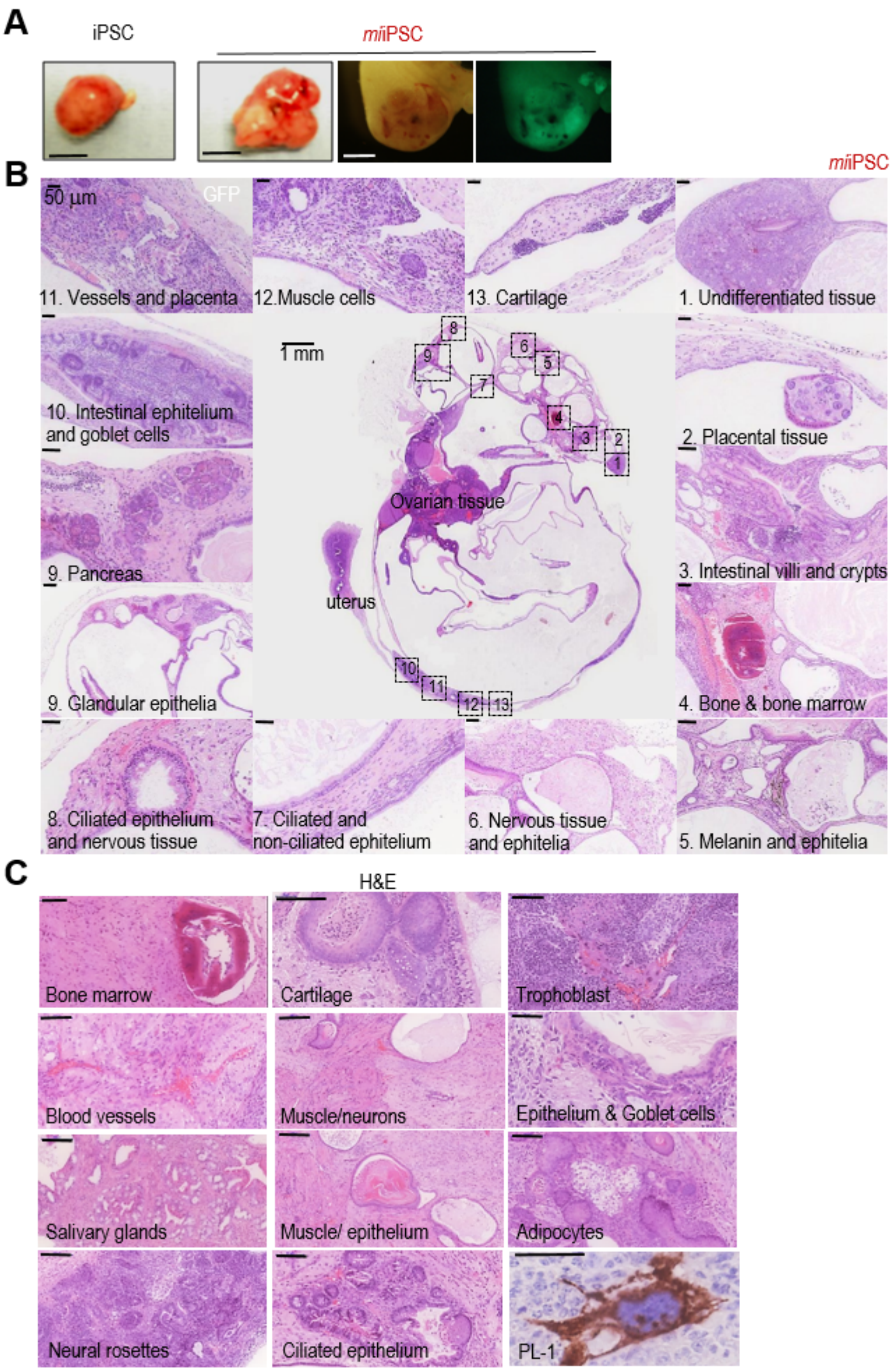

**Figure EV2** Histopahological analysis of teratomas generated from *mi*iPSCs. **A**, Representative images of teratomas 20-25 days after subcutaneous injection of wild-type iPSCs or *mi*iPSCs expressing GFP, as indicated in Fig 2A. Images on the right show an example of a *mi*iPSC-derived embryo-like structure, analyzed 20 days after intraperitoneal (i.p.) injection of *mi*iPSCs expressing GFP. Scale bar, 5 mm (two left panels), 1 mm (two right panels). **B**, Representative example of a highly differentiated teratoma generated from *mi*iPSCs after i.p. injection in nude mice. Most of these complex teratomas were detected in the proximity of the uterus or as ovarian cysts in the host mice. The panel shows higher magnifications (H&E staining) of several differentiated tissues and cells observed in the teratoma. Scale bars, 1 mm (central image) or 50 μm (insets). **C**, Histopathological examples (H&E staining) of specific tissues found in *mi*iPSCs-derived teratomas. Scale bars, 100 μm. A magnification of a trophoblast giant cell stained against Placental Lactogen-1 (PL-1) is also shown (scale bar, 50 μm).

**A**

| | Injected embryos | Tranferred embryos | Recovered embryos | Implantation sites |
|---|---|---|---|---|
| Control hiPSCs P29 | 20 | 14 | 3 | 9 |
| Human *mi*iPSCs P29+0 | 21 | 16 | 8 | 4 |
| Human *mi*iPSCs P36+2 | 21 | 16 | 9 | 5 |

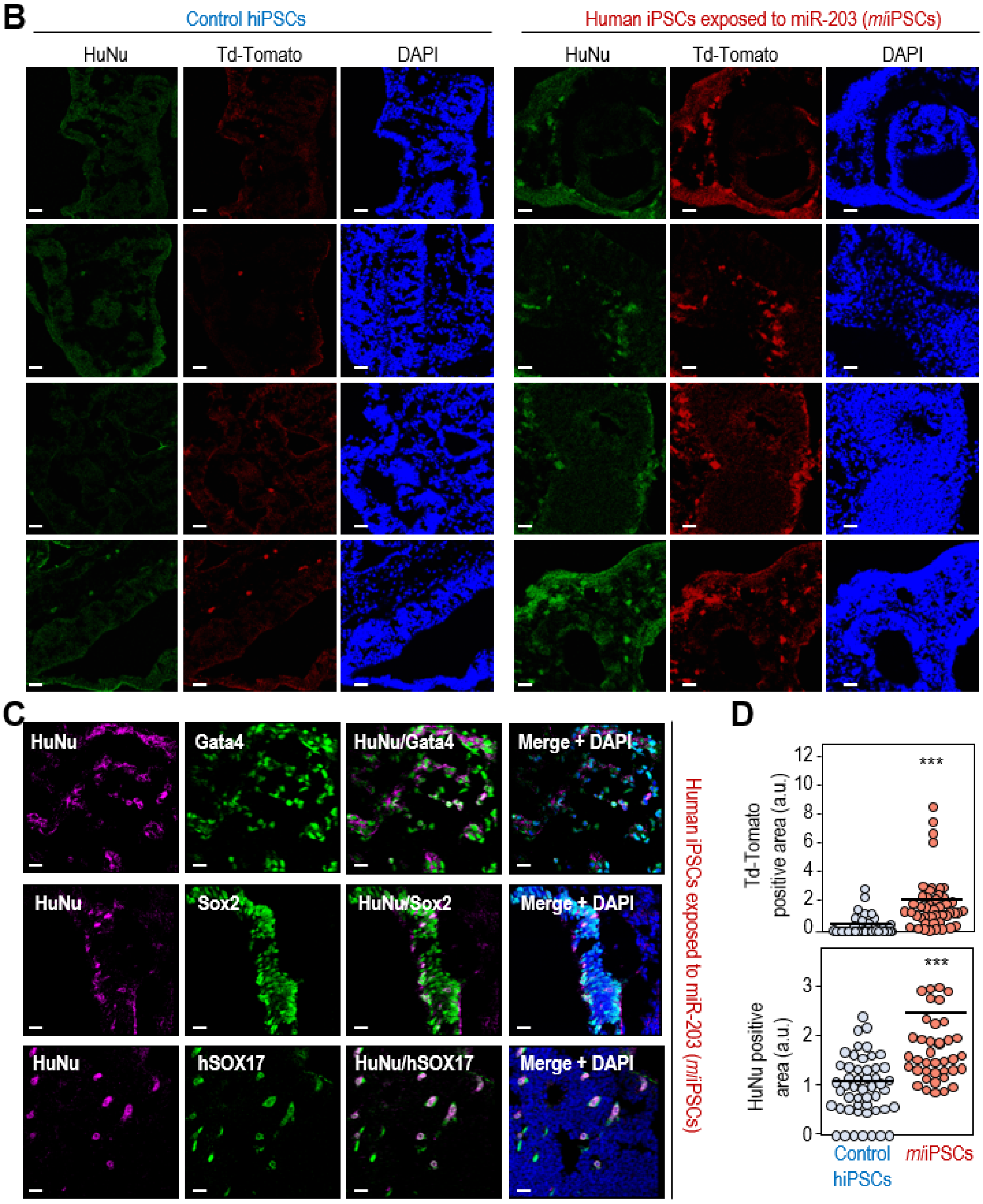

**Figure EV3** miR-203-exposed human iPSCs efficiently contribute to human-mouse interspecies chimeras. A, Summary of interspecies chimera assays in which one single Td-Tomato fluorescent-labeled human iPSC, either control or transiently transfected with miR-203-expressing vectors (*mi*iPSCs), was injected into 8C-stage mouse embryos. These embryos were cultured in vitro for 48 h to reach the blastocyst stage and then transfected to 2.5 day post coitum pseudo-pregnant females. The post-implantation mouse conceptuses were dissected at the E9.5 developmental stage and analyzed by immunofluorescence. B, Representative immunofluorescence images from these E9.5 embryos. Anti-human HuNu antibody was co-stained with Td-Tomato direct fluorescence to detect human cells in E9.5 mouse embryos. Scale bars, 20 μm. C, Representative immunofluorescence images from these E9.5 embryos co-stained with human HuNu antibody and Gata4, Sox2 or hSOX17 antibodies showing differentiation and proper integration of human cells into the mouse embryo. Scale bars, 20 μm. D, Quantification of the Td-Tomato-(d) or HuNu-(e) positive area in cryosections of E10.5 embryos resulting from chimera assays in which 15 Td-Tomato fluorescent-labeled human iPSCs, either control or transiently transfected with miR-203-expressing vectors (*mi*iPSCs) were injected into 8C-stage mouse embryos and their contribution analyzed 48-60 h later. ****P*<0.001 (Student´s t-test).

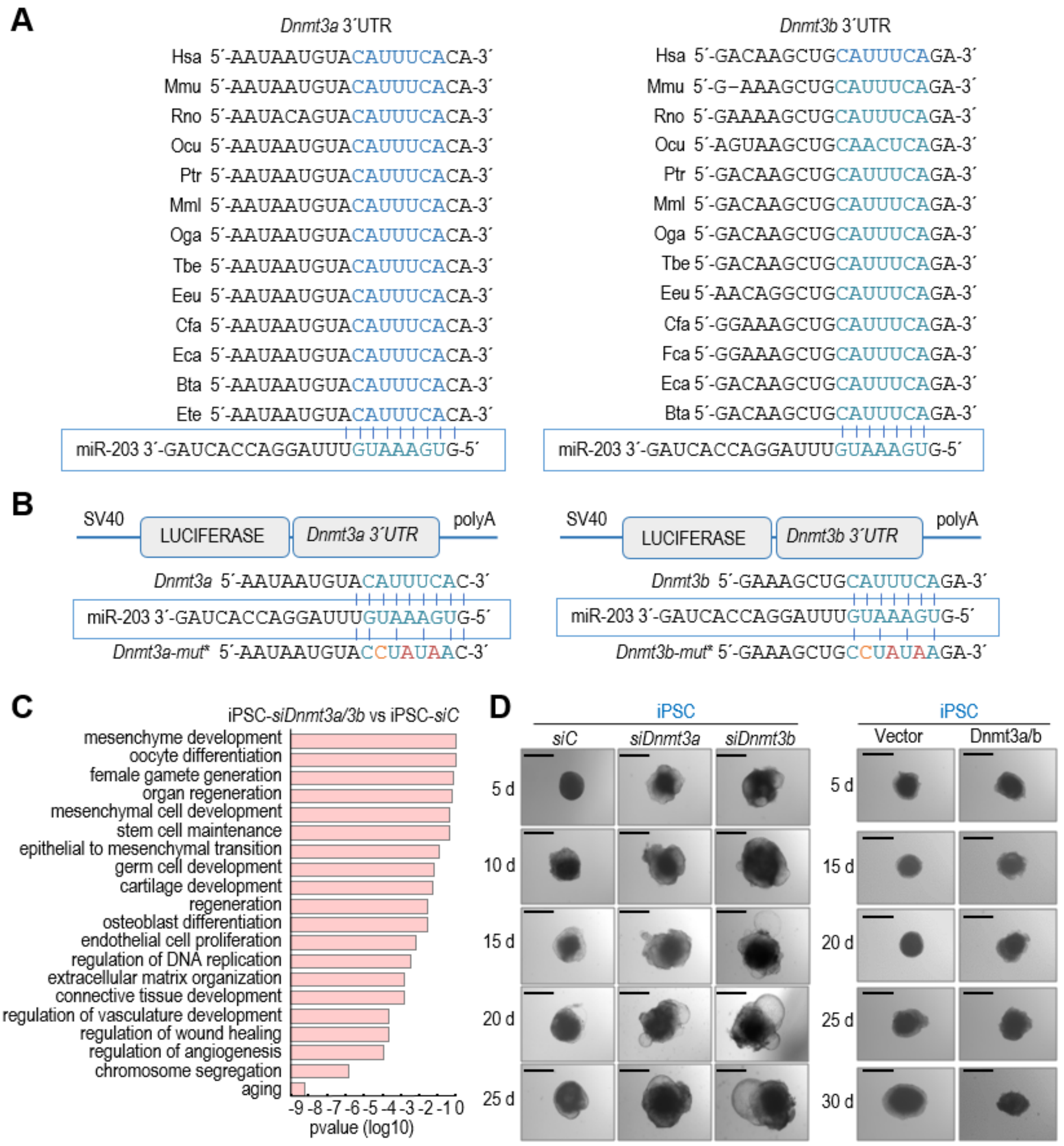

**Figure EV4** miR-203 targets *Dnmt3a/b* 3'-UTR sequences. **A**, *Dnmt3a* and *Dnmt3b* 3´UTR alignment in several representative species. The seed region of the miR-203 target site contained in these 3´-UTRs is highlighted in blue and aligned with the corresponding miR-203 seed sequence. **B**, Schematic representation of the luciferase reporter, carrying the wild-type *Dnmt3a* (left) or *Dnmt3b* (right) complete 3´-UTRs or the corresponding mutated versions, downstream of the luciferase gene. The mutated residues are indicated in red. **C**, Major pathways from the Gene Ontology database upregulated in wild-type iPSCs treated with specific siRNAs against *Dnmt3a* and *Dnmt3b* versus scrambled sequences (siC). **D**, Representative images of embryoid bodies (EBs) generated from wild-type iPSCs treated with specific siRNAs either against *Dnmt3a* or *Dnmt3b* (left panel) or transfected with a combination of miR-203-resistant Dnmt3a/b cDNAs (right panel) at the indicated time points during the differentiation process.

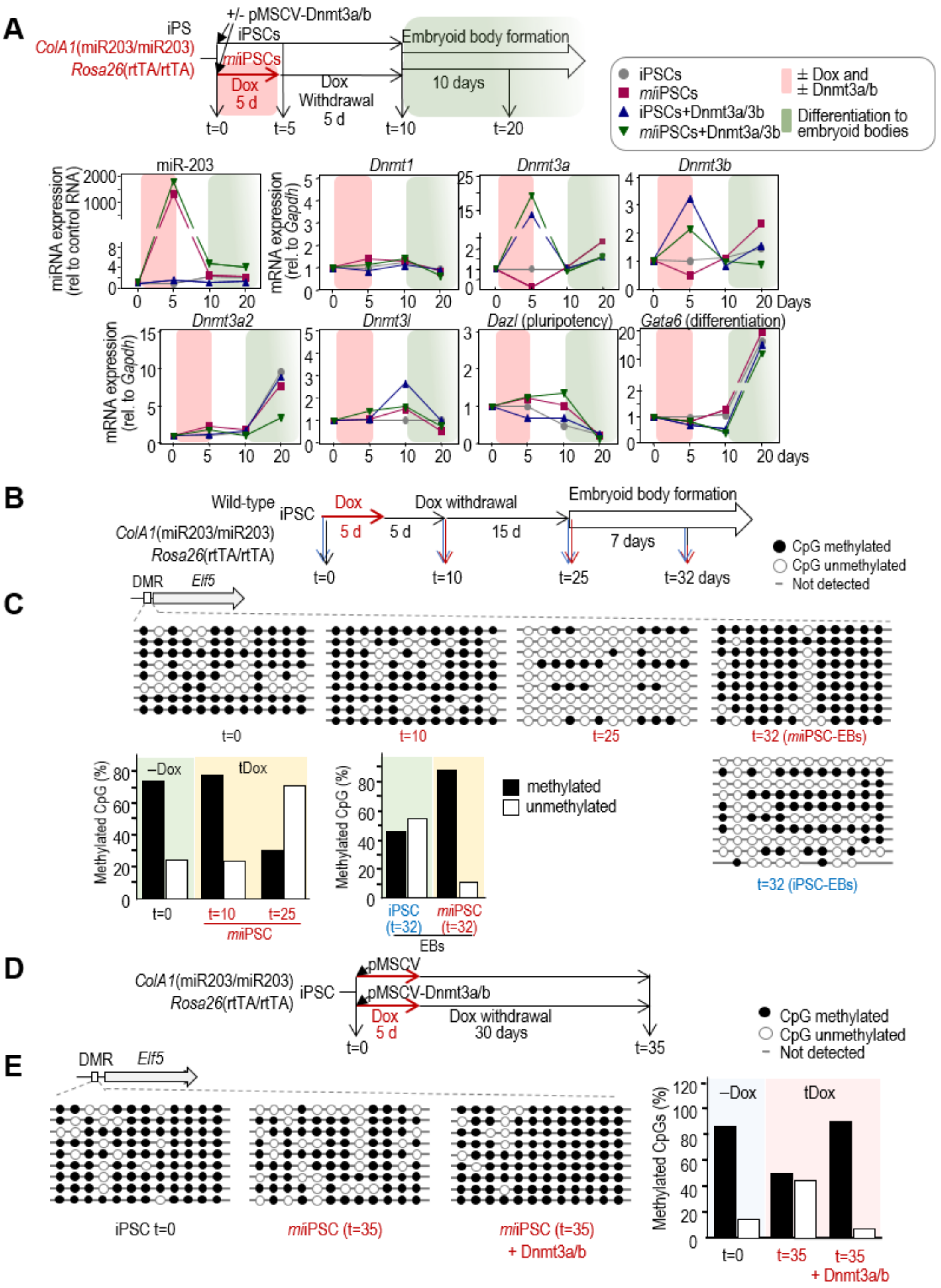

**Figure EV5** Genome-wide methylation of iPSCs or embryoid bodies after transient exposure to miR-203. A, Expression levels, as determined by quantitative PCR, of *miR-203*, and transcripts for the DNA methyltransferases *Dnmt1, Dnmt3a, Dnmt3b, Dnmt3a2* and *Dnmt3l*, or pluripotency (*Dazl*) and differentiation (*Gata6*) markers. *ColA1*(miR-203/miR-203); *Rosa26*(rtTA/rtTA) iPSCs treated or not with doxycycline (Dox) and simultaneously transduced with Dnmt3a/b cDNAs or empty vector were used as shown in the schematic representation of the experimental design. The pink shadow indicates the time-lapse in which the cells were treated or not with Dox and transduced or not with Dnmt3a/b cDNAs. The orange shadow indicates the differentiation process to embryoid bodies. Data are represented as mean of three technical replicates per experiment (n=2 independent experiments). B, Experimental design used for the validation of methylation data in the indicated *Elf5* differentially methylated region (DMR). C, DNA was isolated as indicated and sequenced after bisulfite modification. Eight to ten independent clones were sequenced per condition. Histograms show the percentage of DNA methylation at the *Elf5* DMR in the different conditions. D, Experimental protocol followed to test DNA methylation rescue by miR-203-resistant Dnmt3a/b cDNAs. E, The specific differentially methylated regions (DMR) at the *Elf5* locus were analyzed by PCR amplification and sequencing of bisulphite-modified DNA. The quantification of methylated vs. unmethylated CpGs is shown in the histogram.

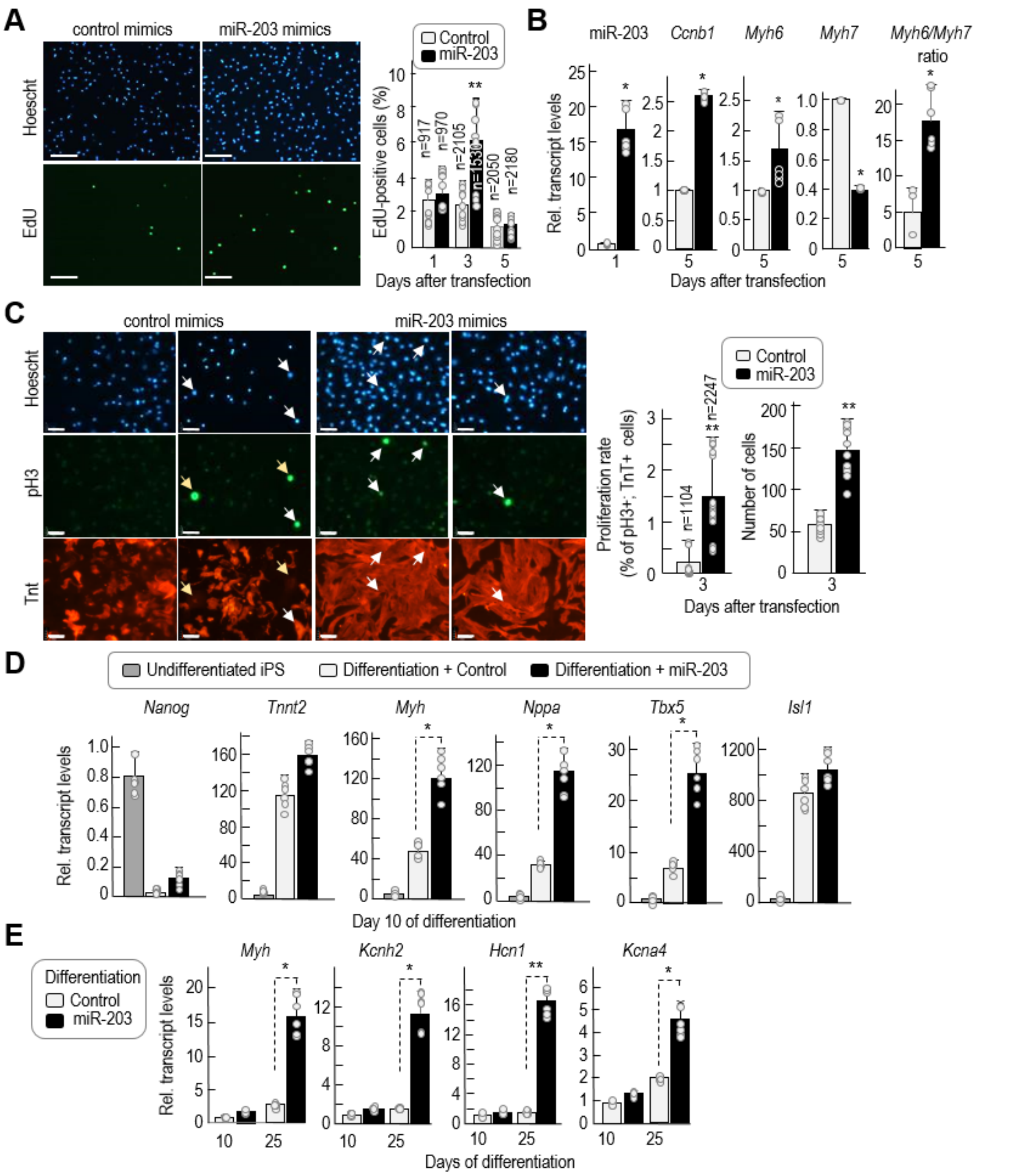

**Figure EV6** Improved cardiomyocyte differentiation and maturation after transient expression of miR-203. A, Representative immunofluorescence showing EdU (green) and nuclei (DAPI, blue) staining of primary cardiomyocytes extracted at postnatal day 1 and transiently transfected with control or miR-203 mimics 24 hours after extraction. Pictures were taken three days after transfection. Scale bar, 60 μm. The histogram shows the percentage of EdU-positive cells at different days post-transfection. Data are mean ± s.d. (n=2 independent experiments with 6 replicates each). B, RNA expression as determined by quantitative PCR of miR-203 (24 hours after transfection) and *Ccnb1*, *Myh6* and *Myh7* transcripts (5 days after transfection). The *Myh6/Myh7* ratio is calculated as an indicator of cardiomyocyte maturation. Data are mean ± s.d. (n=3 independent experiments). C, Representative immunofluorescence showing phospho-Histone 3 (Ser-10; pH3) in green, cardiac Troponin T (cTnT) in red and Hoescht for nuclei staining in blue, in primary cardiomyocytes extracted at postnatal day 1 and transiently transfected with control or miR-203 mimics 24h after extraction. Images were taken three days after transfection. Scale bars, 64 μm. White arrows point to cardiomyocytes positive for pH3. Middle histogram shows the proliferation rate measured as the percentage of pH3 positive cells respect to the total number of cTnT positive cells at day 3 post-transfection. Data are mean ± s.d. (n=2 independent experiments). The plot on the right shows the total number of cells at day 3 post-transfection. Data are represented as mean ± s.d. (n=2 independent experiments). **D**, mRNA levels as determined by quantitative PCR of the indicated transcripts at different time points before and during cardiomyocyte differentiation. iPSCs were transfected either with control mimics or miR-203 mimics, maintained during 15 days in culture and then differentiated *in vitro.* **E**, mRNA levels of the indicated transcripts at different time points during cardiomyocyte differentiation. In **A-E** data are represented as mean ± s.d. (n=2 independent experiments with 6 replicates each). $^{*}P<0.05$, $^{**}P<0.01$ (Student´s *t*-test).

### Supplementary Tables

Supplementary Table 1 | Gene Ontology Analysis of genes upregulated in *mi*iPSCs (5 days in the presence of doxycycline and 30 days after doxycycline withdrawal) versus un-induced iPSCs.

See Excel file.

Supplementary Table 2 | Gene Ontology Analysis of genes significantly deregulated in *mi*iPSC-derived teratomas vs. un-induced iPSC-derived teratomas.

| Gene Ontology | log10 (p-value) | Transcripts |
|---|---|---|
| Extracellular matrix organization (GO:0030198) | -10.9 | *Col1a1, Nid1, Col5a1, Postn, Lox, Col3a1, Lamc1, Nepn, Col1a2, Fn1, Serpinh1, Hspg2, Cyr61, Adamts2, Col11a1, Col5a2, Foxf1a, Itga8, Klf4, Pdgfra, Vmo1* |
| Cell-substrate adhesion (GO:0031589) | -10.0 | *Col1a1, Nid1, Thbs1, Col3a1, Lamc1, Ctgf, Fn1, Sox17, Tek, Cyr61, Fbln2, Itgbl1, Klf4, Nid2, Rell2* |
| Organ morphogenesis (GO:0009887) | -8.4 | *Col1a1, Igf2, Lama1, Thbs1, Col5a1, Hand1, Afp, Gata4, Myl3, Ctgf, Fst, Col4a2, Col1a2, Phlda2, Figf, Snai1, Tgfbr3, Hand2, Shank3, Sox17, Tek, Mybpc3, Hspg2, Cyr61, Enpep, Cited1, Edn1, Actc1, Ahsg, Alx3, Ang, Bmp4, Ccl2, Cfc1, Col11a1, Col5a2, Cryaa, Crygs, Fgf3, Flt1, Foxf1a, Ghr, Hoxa13, Hoxc11, Hoxd9, Itga4, Itga8, Klf4, Mmp2, Msx2, Myh6, Myl2, Pdgfra, Rbp4, Tbx20, Tbxa2r, Wnt6* |
| Cell adhesion (GO:0007155) | -7.4 | *Col1a1, Lama1, Nid1, Thbs1, Col5a1, Postn, Col3a1, Lamc1, Ctgf, Svep1, Wisp1, Fn1, Pcdh11x, Sox17, Spon2, Tek, Mybpc3, Hspg2, Cyr61, Col12a1, Lamc3, Podxl, Adam12, B4galnt2, Cdh3, Cntnap1, Cntnap5b, Col11a1, Col6a1, Col9a1, Dpp4, Fbln2, Hapln1, Igsf9b, Itga4, Itga8, Itgbl1, Kcp, Klf4, Ncan, Nid2, Pcdh10, Pcdh19, Rell2, Vnn1* |
| Tissue development (GO:0009888) | -6.1 | *Col1a1, Lama1, Col5a1, Gata6, Postn, Hand1, Col3a1, Gata4, Myl3, Ctgf, Fst, Col1a2, Snai1, Pth1r, Tgfbr3, Hand2, Igfbp1, Shank3, Sox17, Mybpc3, Cyp26a1, Hspg2, Adamts2, Edn1, Actc1, Ahsg, Bmp4, Cdkn2a, Cfc1, Col11a1, Col5a2, Col9a1, Crygs, Foxf1a, Ghr, Ghsr, Hoxd9, Klf4, Myh6, Myl2, Ntf3, Pitx1, Pln, Rbp4, Sprr2k, Sprr4, Srpk3, Tcf23, Trpv1, Unc45b, Upk3a* |
| Embryonic skeletal system development (GO:0048706) | -5.4 | *Col1a1, Hspg2, Alx3, Col11a1, Hoxb9, Hoxd9, Klf4, Pcsk5, Rbp4* |
| Skeletal system development (GO:0001501) | -5.1 | *Col1a1, Igf2, Col3a1, Ctgf, Col1a2, Pth1r, Hspg2, Edn1, Ahsg, Alx3, Bmp4, Casr, Col11a1, Col5a2, Col9a1, Ghr, Hoxb9, Hoxc11, Hoxd9, Klf4, Mmp2, Nppc, Pcsk5, Pdgfra, Pitx1, Rbp4, Twist2* |
| Chordate embryonic development (GO:0043009) | -4.9 | *Col1a1, Gata6, Hand1, Gata4, Tgfbr3, Hand2, Sox17, Hspg2, Cited1, Edn1, Serpina1b, Cdx2, Alx3, Apba1, Bmp4, Col11a1, Dab2, Foxf1a, Hoxb9, Hoxd9, Insl3, Klf1, Klf4, Myh6, Pcsk5, Pdgfra, Rbp4* |
| Embryonic development ending in birth or egg hatching (GO:0009792) | -4.8 | *Col1a1, Gata6, Hand1, Gata4, Tgfbr3, Hand2, Sox17, Hspg2, Cited1, Edn1, Serpina1b, Cdx2, Alx3, Apba1, Bmp4, Col11a1, Dab2, Evx1, Foxf1a, Hoxb9, Hoxd9, Insl3, Klf1, Klf4, Myh6, Pcsk5, Pdgfra, Rbp4* |
| Blood vessel development (GO:0001568) | -4.7 | *Col1a1, Thbs1, Col5a1, Hand1, Lox, Col3a1, Ctgf, Col4a2, Col1a2, Figf, Tgfbr3, Hand2, Sox17, Tek, Cyr61, Enpep, Cited1, Edn1, Cdx2, Ang, Bmp4, Flt1, Foxf1a, Gja4, Itga4, Klf4, Mmp2, Tbx20, Tbxa2r* |
| Vasculature development | -4.7 | *Col1a1, Thbs1, Col5a1, Hand1, Lox, Col3a1, Ctgf, Col4a2, Col1a2, Figf, Tgfbr3, Hand2, Sox17, Tek, Cyr61, Enpep, Cited1, Edn1, Cdx2, Ang, Bmp4, Flt1, Foxf1a,* |

| | | |
|---|---|---|
| (GO:0001944) | | *Gja4, Itga4, Klf4, Mmp2, Tbx20, Tbxa2r* |
| Skeletal system morphogenesis (GO:0048705) | -4.6 | *Col1a1, Ctgf, Hspg2, Alx3, Col11a1, Ghr, Hoxd9, Klf4, Mmp2, Pdgfra* |
| Regulation of cell adhesion (GO:0030155) | -4.5 | *Col1a1, Lama1, Nid1, Thbs1, Cyr61, Podxl, B4galnt2, Dpp4, Fbln2, Klf4, Rell2* |
| Collagen fibril organization (GO:0030199) | -4.4 | *Col1a1, Col5a1, Lox, Col3a1, Col1a2, Serpinh1, Adamts2, Col11a1, Col5a2, Klf4* |
| Embryonic development (GO:0009790) | -4.1 | *Col1a1, Lama1, Gata6, Hand1, Gata4, Tgfbr3, Hand2, Shank3, Sox17, Hspg2, Cited1, Edn1, Serpina1b, Cdx2, Alx3, Apba1, Bmp4, Cfc1, Col11a1, Dab2, Evx1, Foxf1a, Hoxa13, Hoxb9, Hoxc11, Hoxd9, Insl3, Itga8, Klf1, Klf4, Msx2, Myh6, Pcsk5, Pdgfra, Rbp4, Tbx20, Thbd* |
| Skin development (GO:0043588) | -4.0 | *Col1a1, Col5a1, Col3a1, Col1a2, Adamts2, Col5a2, Klf4* |
| Positive regulation of cell-substrate adhesion (GO:0010811) | -3.9 | *Nid1, Thbs1, Cyr61, Fbln2, Klf4, Rell2* |
| Cell-matrix adhesion (GO:0007160) | -3.8 | *Nid1, Col3a1, Ctgf, Fn1, Sox17, Tek, Itgbl1, Klf4, Nid2* |
| Cell development (GO:0048468) | -3.6 | *Lamc1, Prdm6, Pth1r, Tgfbr3, Hand2, Sox17, Gdf9, Mybpc3, Hba-x, Cyp26a1, Edn1, Cdx2, Actc1, Bmp4, Col11a1, Cryaa, D3Bwg0562e, Dab2, Evx1, Grin2a, Hoxd9, Insl3, Klf1, Klf4, Lgi4, Lmx1a, Myh6, Myl2, Ntf3, Pde3a, Pdgfra, Pitx1, Rbp4, Rtn4rl1* |
| Collagen biosynthetic process (GO:0032964) | -3.4 | *Col1a1, Col5a1, Serpinh1, F2* |
| Heart development (GO:0007507) | -3.2 | *Col5a1, Gata6, Hand1, Col3a1, Gata4, Myl3, Tgfbr3, Hand2, Sox17, Mybpc3, Hspg2, Edn1, Actc1, Bmp4, Cfc1, Col11a1, Itga4, Klf4, Msx2, Myh6, Myl2, Pcsk5, Pln, Rbp4, Tbx20* |
| Behavior (GO:0007610) | -3.2 | *Lama1, Thbs1, Lamc1, Alb, Slc6a3, Sox17, Cyr61, Gabra5, a, Adcy1, Apba1, Apln, Aplnr, Casr, Ccl2, Ccl7, Chrna4, Cyp11a1, Fpr1, Ghsr, Gpr34, Grin2a, Hoxd9, Itga8, Klf4, Trpv1* |
| Anatomical structure formation involved in morphogenesis (GO:0048646) | -3.1 | *Col1a1, Thbs1, Hand1, Ctgf, Col4a2, Figf, Tgfbr3, Hand2, Shank3, Sox17, Tek, Mybpc3, Cyr61, Enpep, Edn1, Cdx2, Actc1, Ang, Bmp4, Fgf3, Flt1, Hoxc11, Klf4, Mmp2, Myh6, Myl2, Rbp4, Tbx20, Tbxa2r* |

Supplementary Table 3 | List of miR-203 predicted targets among the transcripts downregulated in *mi*iPSCs and involved in the epigenetic regulation of gene expression (GO0040029).

| Transcript | Predicted miR-203 targets | | |
|---|---|---|---|
| | *mi*iPSC vs iPSC log2(fold_change) | Agreement[a] | Computational Predictions (method\|target-site\|start\|stop\|score) |
| *Dnmt3a* | -0.53 | 1.43 | Miranda\|7mer-m8\|5437\|5458\|1.42576278100035\| |
| *Dnmt3b* | -2.36 | 1.33 | Miranda\|Offset 3-8 6mer\|612\|634\|1.32739957876685\| |
| *Uhrf2* | -0.52 | 1.31 | Miranda\|N/A\|790\|809\|1.31334769273349\| |
| *Trim27* | -0.74 | 1.29 | Miranda\|Offset 3-8 6mer\|87\|107\|1.28524392066678\| |
| *Tet1* | -1.05 | 1.29 | Miranda\|8mer\|128\|149\|1.28524392066678\| |
| *Apobec1* | -0.67 | 1.26 | Miranda\|N/A\|679\|700\|1.25714014860006\| |
| *Ctcf* | -0.65 | 1.24 | Miranda\|6mer\|925\|947\|1.24308826256671\| |
| *Hells* | -1.40 | 1.24 | Miranda\|7mer-m8\|2779\|2799\|1.24308826256671\| |
| *Klf2* | -1.11 | 1.23 | Miranda\|N/A\|414\|437\|1.22903637653335\| |
| *Mier1* | -0.86 | 1.21 | Miranda\|6mer\|2450\|2471\|1.21498449049999\| |
| *Smarca5* | -0.63 | 1.20 | Miranda\|N/A\|1405\|1426\|1.20093260446663\| |
| *Dnd1* | -1.34 | 1.20 | Miranda\|Offset 3-8 6mer\|247\|267\|1.20093260446663\| |
| *Brca1* | -1.51 | 1.20 | Miranda\|6mer\|692\|713\|1.20093260446663\| |
| *Dpy30* | -0.90 | 0.27 | Miranda\|N/A\|98\|118\|1.25714014860006\| Rnahybrid\|Offset 1-7 8mer\|109\|117\|-0.759910346569166\| |
| *Rlim* | -0.51 | -0.75 | Pita\|8mer\|520\|526\|-0.752279554170068\| |
| *Rbm3* | -1.04 | -0.76 | Pita\|8mer\|442\|448\|-0.756298393575608\| |
| *H2afy* | -0.54 | -0.76 | Targetscan\|7mer-m8\|474\|480\|-0.764286890785572\| |

[a] The agreement score of the target, which is a weighted mean (using each method's experimental AUC) of the computational methods scores in the prediction group.

Supplementary Table 4 | Differentially methylated positions (DMP) in *mi*iPSC (t=10 versus t=0 and t=25 versus t=0)

See Excel file.

Supplementary Table 5 | Extended data of genome-wide methylation analysis.

See Excel file.

Supplementary Table 6 | List of genes included in the 2-cell-embryo signature (Biase et al, 2014) and their TSS methylation level in *mi*iPSCs.

See Excel file.

Supplementary Table 7 | A comparison between miPSCs and other PSCs previously published in representative studies.

See Excel file.

Supplementary Table 8 | Antibodies used in this work.

| Antibodies | | | | |
|---|---|---|---|---|
| Antigen | Catalog Number | Clone number | Source Ig | Source |
| AFP | AF5369 | - | goat | R&D Systems |
| Cd34 | Ab8158 | MEC14.7 | rat | Abcam |
| CK8 | AM-TROMA I | - | mouse | CNIO Monoclonal Antibodies Core Unit |
| Gata4 (C-20) | Sc-1237 | - | goat | Santa Cruz Biotechnology |
| GFP | 11 814 460 001 | 7.1+13.1 | mouse | Roche |
| Histone 3 (phospho-Ser10) | 06-570 | - | rabbit | Millipore |
| HuNu | NBP2-34342 | - | mouse | Novus |
| Oct4 | Ab19857 | - | rabbit | Abcam |
| Oct4 | C30A3 | - | rabbit | Cell Signalling |
| PL-1 | Sc-34713 | - | goat | Santa Cruz Biotechnology |
| Sox17 | AF1924 | - | human | R&D Systems |
| Sox2 | AF2018 | - | h/mouse/rat | R&D Systems |
| Sox2 | 3728 | C70B1 | rabbit | Cell Signaling Technology |
| Ter119 (LY-76) | 550565 | Ter119 | rat | BD Bioscience |
| Troponin T | Ab8295 | 1C11 | mouse | Abcam |

Supplementary Table 9 | Oligonucleotides used in this work.

| Gene | Forward Oligonucleotide (5'-3) | Reverse Oligonucleotide (5'-3) |
|---|---|---|
| *qRT-PCR (mouse genes)* | | |
| *Dazl* | GGTTTTACCACCCGAACTCTG | TGTGGTTGCTGATGAAGACTG |
| *Dnmt1* | CAGAGACTCCCGAGGACAGA | TTTACGTGTCGTTTTTCGTCTC |
| *Dnmt3a* | AAACGGAAACGGGATGAGT | ACTGCAATTACCTTGGCTTTCT |
| *Dnmt3a2* | GGGCAAACTGAAGTAGTGATGA | TTACACGGCACCTGCTGA |
| *Dnmt3b* | CCCTCCCCCATCCATAGT | TCTGCTGTCTCCCTTCATTGT |
| *Dnmt3l* | AAGTGAACCGACGGAGCAT | CCGAGTGTACACCTGGAGAGTT |
| *Ecat1* | TGTGGGGCCCTGAAAGGCGAGCTGAGAT | ATGGGCCGCCATACGACGACGCTCAACT |
| *Eras* | ACTGCCCCTCATCAGACTGCTACT | CACTGCCTTGTACTCGGGTAGCTG |
| *Esg1* | GAAGTCTGGTTCCTTGGCAGGATG | ACTCGATACACTGGCCTAGC |
| *Fgf4* | CGTGGTGAGCATCTTCGGAGTGG | CCTTCTTGGTCCGCCCGTTCTTA |
| *Gapdh* | AGGTCGGTGTGAACGGATTTG | TGTAGACCATGTAGTTGAGGTCA |
| *Gata6* | ACCTTATGGCGTAGAAATGCTGAGGGTG | CTGAATACTTGAGGTCACTGTTCTCGGG |
| *Gdf3* | GTTCCAACCTGTGCCTCGCGTCTT | AGCGAGGCATGGAGAGAGCGGAGCAG |
| *Hcn1* | TGAAGCTGACAGATGGCTCTT | CTGGCAGTACGACGTCCTTT |
| *Isl1* | TTGTACGGGATCAAATGCGCCAAG | AGGCCACACAGCGGAAACA |
| *Kcna4* | TCATTGCTCTGACCTGATGC | TCACTCAGCTCCCTCAGGAT |
| *Kcnh2* | ACGCTTACTGCCAGGGTGAC | GCCGACTGGCAACCAGAG |
| *Myh* | CTCAAGCTCATGGCCACTCT | GCCTCCTTTGCTTTTACCACT |
| *Nanog* | CAGGTGTTTGAGGGTAGCTC | CGGTTCATCATGGTACAGTC |
| *Nppa* | GAACCAGAGGGGAGAGACAGAG | CCCTCAGCTTGCTTTTTAGGAG |
| *Tbx5* | AAATGAAACCCAGCATAGGAGCTGGC | ACACTCAGCCTCACATCTTACCCT |
| *Tnnt2* | GGCAGCGGAAGAGGATGCTGAA | GAGGCACCAAGTTGGGCATGAACGA |
| *qRT-PCR (rat genes)* | | |
| *Ccnb1* | GGAGATGAAGATTCTGAGAGTTCTG | GTATGCTGCTCCACATCGAC |
| *Gapdh* | GGCAAGTTCAATGGCACAGT | TGGTGAAGACGCCAGTAGACTC |
| *Myh6* | GGGCTGGAGCACTGAGAG | GAGAGAGGAACAGGCAGGAA |
| *Myh7* | ATGGCGGATCGAGAGATG | GGTCAAAGGGCCTGGTCT |
| *DNA methylation analysis (mouse genes)* | | |
| *Elf5 (a)* | TAAAGGTTGTAATGAATAGATATTAGGTT | AACTACTTACTTAAAAACAAATAATAACTAAA |
| *Elf5 (b)* | TAAAGGTTGTAATGAATAGATATTAGGTT | AAATAATAACTAAATCCAAACAAAAAA |
| *Sirt6* | TTTGGTTTTTTTTAGGTTATGTTAGGATTT | CACTTACCTCTACCTCCCAATAAAAAA |
